# Excess BAFF Alters NR4As Expression Levels and Breg Function of Human Precursor-like Marginal Zone B-cells in the Context of HIV-1 infection

**DOI:** 10.1101/2022.08.15.504021

**Authors:** Kim Doyon-Laliberté, Matheus Aranguren, Michelle Byrns, Josiane Chagnon-Choquet, Matteo Paniconi, Jean-Pierre Routy, Cécile Tremblay, Marie-Claude Quintal, Nathalie Brassard, Daniel E Kaufmann, Johanne Poudrier, Michel Roger

**Affiliations:** Centre de Recherche du Centre Hospitalier de l’Université de Montréal (CRCHUM), Montréal, Qc, Canada; Département de Microbiologie, Infectiologie et Immunologie de l‘Université de Montréal, Montréal, Qc, Canada; Service d’Aide à la Formation Interdisciplinaire et à la Réussite Étudiante (SAFIRE), Faculté des Arts et Sciences de l’Université de Montréal, Montréal, Qc, Canada; Department of Medicine, McGill University Health Centre, McGill University, Montréal, Qc, Canada; Centre Hospitalier Ste-Justine de l’Université de Montréal, Montréal, Qc, Canada; Département de Médecine de l‘Université de Montréal, Montréal, Qc, Canada

**Author notes:** These authors have contributed equally to this work and share first authorship (Kim Doyon-Laliberté, Matheus Aranguren). These authors have contributed equally to this work and share senior authorship (Johanne Poudrier, Michel Roger). **Correspondence:** Johanne Poudrier PhD and Michel Roger MD PhD, CRCHUM, Tour Viger 900 rue St-Denis, Montréal, Canada H2X 0A9.

**Keywords:** HIV, Breg, NR4A, MZp, BAFF

## Abstract

We have shown that excess B-cell activating factor (BAFF) in the blood of HIV-infected individuals, is concomitant with increased frequencies of precursor-like marginal zone (MZp) B-cells, early on and despite successful antiretroviral therapy (ART). We have recently reported that in HIV-uninfected individuals, MZp possess a strong B-cell regulatory (Breg) potential. As such, MZp B-cells highly express IL-10, the orphan nuclear receptors (NR)4A1, NR4A2, NR4A3, the regulatory molecule CD83, as well as ectonucleotidases CD39 and CD73, all of which are associated with regulation of inflammation. Moreover, the Breg function of MZp B-cells involves CD83 signals. Herein, in order to address the impact of HIV infection and excessive BAFF environment on MZp B-cells and their regulatory capacities, we have performed transcriptomic analyses by RNA-seq of sorted MZp B-cells from the blood of HIV-infected progressors. The Breg profile and function of blood MZp B-cells from HIV-infected progressors were assessed by flow-cytometry and light microscopy high content screening (HCS) analyses, respectively. In addition, the effects of excess BAFF on the Breg profile of MZp B-cells from HIV-uninfected controls were investigated *in vitro*. We report significant downregulation of NR4A1, NR4A2, NR4A3 and CD83 gene transcripts in blood MZp B-cells from HIV-infected progressors when compared to HIV-uninfected controls. NR4A1, NR4A3 and CD83 protein expression levels and Breg function were also downregulated in blood MZp B-cells from HIV-infected progressors and not restored by ART. Moreover, we observe decreased expression levels of NR4A1, NR4A3, CD83 and IL-10 by blood and tonsillar MZp B-cells from HIV-uninfected individuals following treatment with excess BAFF, which significantly diminished their regulatory function. These findings suggest that excess BAFF contributes to the alteration of the Breg potential of MZp B-cells, which could lead to a loss of “immune surveillance”, during HIV infection and possibly in other situations where BAFF is found in excess.

**Author Summary:** The precursor-like marginal zone (MZp) B-cell population, we previously described in human blood and tonsils, presents with an important regulatory “Breg” potential, depicted by elevated nuclear receptor (NR)4As expression levels, similarly to Tregs, and to our knowledge currently underexplored in human Breg studies. Herein, we present the impact that a chronic inflammatory context such as HIV-infection, and its excessive B-cell activating factor (BAFF) environment, may exert on the Breg capacities of MZp, both *ex vivo* and *in vitro*, significantly affecting their NR4As expression levels and Breg function. These findings are of growing significance, especially with the recently described importance of MZ B-cell NR4A1 expression in atherosclerosis immune surveillance. The finding that immune surveillance may be altered in circumstances of chronic inflammation and excessive BAFF, is of pivotal interest, as treated HIV-infected individuals often prematurely develop co-morbidities associated with aging such as cardiovascular diseases (CVD). Moreover, excess BAFF has been reported in several inflammatory autoimmune contexts where CVD is the leading cause of death.

## Introduction

Currently, over 37 million individuals are living with the human immunodeficiency virus (HIV) [1]. Although over 66% of these individuals are successfully treated with antiretroviral therapy (ART), inflammation persists [1, 2]. Consistently, we have previously reported that expression levels of the tumor necrosis factor (TNF) family member B-cell Activating Factor (BAFF) are in excess in the blood of HIV-infected progressors from the Montreal Primary HIV Infection (PHI) cohort. This upregulation is already present in the acute phase of infection and is still observed 9-12 months post-ART [3]. Moreover, our recent studies suggest that excess BAFF perseveres in HIV-infected individuals on ART for more than 15 years of the Canadian HIV and Aging Cohort Study (CHACS) (Aranguren *et al; unpublished observations*). Excess BAFF is associated with viral factors, such as Nef, and non-viral factors, such as elements of microbial translocation, and correlates with hypergammaglobulinemia and deregulation of the B-cell compartment [4–6]. BAFF is an important growth factor that helps shape the marginal zone (MZ) B-cell pool. As such, we found that excess BAFF is associated with increased frequencies of a CD19^+^CD1c^+^CD21^lo^IgM^hi^CD27^+^CD10^+^ B-cell population, sharing features of transitional immature and MZ B-cells, we designated “Precursor-like” Marginal Zone (MZp) B-cells, in the blood of PHI HIV-infected progressors [3], and more recently in the blood of CHACS participants (Aranguren et al; *unpublished observations*). We have reported similar observations with HIV-Transgenic (Tg) mice [7], SIV-infected macaques [8] and with HIV-infected Beninese commercial sex workers [9, 10], suggesting that excess BAFF and deregulated MZp B-cells are reliable markers of inflammation and disease progression in the context of HIV.

BAFF binds to three receptors among B-cell subsets: BAFF-R, transmembrane activator and calcium modulator and cytophilin ligand interactor (TACI) and B-cell maturation antigen (BCMA), the second which is highly expressed by MZ populations [11]. MZ are first-line B-cells that respond quickly to blood borne antigens via their polyreactive BCR and numerous Pattern Recognition Receptors (PRR) such as C-type lectins and Toll-like Receptors (TLR). MZ B-cells have been shown to class switch their immunoglobulins (Ig) from IgM to IgG or IgA in a BAFF-dependant manner via TACI [11]. Importantly, MZ B-cells were shown to bind to the envelope (Env) proteins gp120 and gp41 of HIV, via C-type lectins and TLR10, respectively [12, 13]. Whether Ig produced by MZ B-cells confer some level of protection or contribute to hyperglobulinemia and autoimmune manifestations in excessive BAFF contexts such as with HIV is currently under investigation.

Importantly, we have recently reported that in HIV-uninfected individuals, blood and tonsil MZp B-cells present features associated with strong B-cell regulatory (Breg) capacities [14]. Indeed, transcriptomic and flow-cytometry analyses demonstrated that MZp B-cells express high levels of IL-10, IL-35 and TGFβ, as well as the ectonucleotidases CD39 and CD73. Strikingly, as for regulatory T-cells (Tregs) [15, 16], we found that MZp B-cells highly express the orphan Nuclear Receptors (NR)4A1, NR4A2 and NR4A3, as well as the immunoregulatory molecule CD83 [14], whose expression was shown to be directly modulated by NR4As [17]. Notably, we have shown that the *in vitro* Breg function of tonsil MZp B-cells involved CD83 signaling [14].

NR4As transcription factors are involved in the regulation of the inflammatory response and their expression is increased upon signalling via the TCR or BCR [18], amongst others. They are important regulators of differentiation, proliferation and apoptosis of immune cells [19]. The importance of NR4As in immune regulation is highlighted by their importance for Tregs, as NR4As directly promote and maintain the expression of FoxP3 [16]. Also, increased expression of NR4A1 in myeloid cells is associated with a diminished T-cell activation profile [20]. Furthermore, NR4A1-3 knockout mice develop systemic autoimmune diseases [19, 21].

Given the importance of BAFF for MZ B-cell activity, we have herein assessed the impact of the HIV infection and excess BAFF on the Breg profile and function of MZp B-cells. We show that NR4A1-3 as well as CD83 gene and protein expression levels are severely downregulated in blood MZp B-cells from PHI HIV-infected progressors when compared to HIV-uninfected controls and HIV-elite controllers (EC), whose plasma viral load remain undetectable in absence of ART. Importantly, this downregulation correlates with reduced MZp Breg function. Moreover, this loss of Breg function could be related to the exhaustion profile we describe for this population. We also find that excess BAFF, both as soluble and membrane forms, directly downregulate NR4A1-3 and CD83 expression levels by MZp from the tonsils of HIV-uninfected donors *in vitro*. Total tonsillar membrane BAFF expression levels were also found to correlate with MZp Breg efficiency, as donors expressing relatively low BAFF levels at baseline had a stronger Breg function than donors who had higher BAFF levels at baseline. Strikingly, adding soluble BAFF was found to impede MZp Breg function in low BAFF expressing tonsillar donors. Altogether suggesting that in circumstances of excess BAFF, alteration of MZp Breg profile and function may constitute a major threat to “immune surveillance”. A term utilized by the group of Nus *et al* when describing that depletion of NR4A1 in MZ B-cell populations exacerbated atherosclerosis, thus alluding to the MZ B-cells capacity of “immune surveillance” to prevent atherosclerosis, and possibly other co-morbidities associated with chronic inflammatory conditions [22, 23].

## Materials and Methods

### Specimen collection and clinical data

Cryopreserved blood specimens from 13 male HIV-1 infected classic progressors were selected from the Montreal PHI cohort specimen bank. For simplicity, the term HIV is being used throughout this manuscript. Clinical data from these individuals can be found in Table 1. The date of infection was estimated based on clinical and laboratory results, using criteria established by the Acute HIV Infection and Early Disease Research Program (National Institute of Allergy and Infectious Diseases [NIAID], Bethesda, MD). Blood samples had been collected at two time points: the acute phase of infection (ie: 0-3 months after HIV acquisition) and/or early phase of infection (ie: 5-8 months after HIV acquisition), as well as 96 weeks after HIV acquisition, with an average of 92 weeks of ART treatment (Table 1). Cryopreserved blood samples from 3 male EC were obtained from the Montreal Long Term Non-Progessor (LTNP) cohort [24]. Cryopreserved blood samples from 18 male HIV-negative healthy volunteers were obtained from the Montreal PHI cohort, to be used as HIV-uninfected controls. Plasma viral loads and blood CD4+ T-cell counts were determined as reported previously [25]. None of the subjects had syphilis, hepatitis B, or hepatitis C. Written informed consent was obtained from all subjects, and research conformed to the guidelines of, and was approved by, the University of Montreal Hospital Research Center Ethics Review Board (project reference SL05.028).

**Table 1:** Socio-demographic and data characteristics of the cohorts used for the RNAseq and flow cytometry analyses.

| Group | RNA-seq analysis |  |  | Flow-Cytometry analysis |  |  |
| --- | --- | --- | --- | --- | --- | --- |
|  | HIV+ (n = 3) | EC (n = 3) | p-value | HIV+ ART- (n = 10) | HIV+ ART+ (n = 10) | p-value |
| Age | 38,67 (28 – 44) | 38,67 (37- 40) | 0,7 | NA | NA | NA |
| Viral load | 213353 (51045 – 418465) | NA<sup>1</sup> | NA | 1258139 (374 – 6193724) | 41.4 (20 – 234) | 0,002 |
| CD4+ T-cell count (cells/mm<sup>3</sup>) | 540 (380 – 760) | 819 (737 – 920) | 0,2 | 447 (284 – 590) | 776.5 (565 – 1210) | 0,0006 |
| CD8+ T-cell count (cells/mm<sup>3</sup>) | 871 – (520 – 1153) | 821 (303 - 1130) | >0,99 | 1128 (0 – 3190) | 634.3 (210 – 1080) | 0,06 |
| ART duration (weeks) | NA | NA | NA | NA | 92 (84 – 96) | NA |

Statistical analyses were done between samples selected from HIV-infected progressors (HIV+) and elite controllers (EC), and between HIV+ untreated (ART-) and treated (ART+) individuals. Statistical significance of difference in clinical data between HIV-infected progressors before and after ART was assessed with the Wilcoxon matched-pairs test. Statistical significance of difference between HIV-infected progressors and EC was assessed with the Mann-Whitney U test
<sup>1</sup> Undetectable viral loads were found in the 3 ECs.

### Cell Sorting of Human Blood MZ and MZp B-Cells, RNA Isolation, and Sequencing

Peripheral blood mononuclear cells (PBMCs) from 3 HIV-uninfected controls, 3 HIV-classic progressors at 5-8 months post-infection and 3 EC had been isolated on Ficoll gradients, re-suspended in cryopreservation medium containing 90% heat-inactivated fetal bovine serum (hi-FBS) (Wisent Inc., Montreal, QC, Canada) and 10% dimethyl sulfoxide (DMSO), and stored in liquid nitrogen until use. Cells were thawed, washed in IMDM (Iscove’s Modified Dulbecco’s Medium, Gibco Life Technologies, New York, NY, USA), and processed for cell sorting with a FACSAriaIII apparatus. Live/dead exclusion was performed using LIVE/DEAD Fixable Aqua Dead Cell Stain (Invitrogen Thermo Fisher Scientific, Eugene, OR, USA). Non-specific binding sites were blocked using fluorescence-activated cell sorting (FACS) buffer (1× PBS, 2% hi-FBS) supplemented with 20% hi-FBS and 10 μg mouse IgG (Sigma-Aldrich, St-Louis, MO, USA). Cells were stained using the following conjugated mouse anti-human monoclonal antibodies (mAbs): PacificBlue-anti-CD19, APC-Cy7-anti-CD10 (BioLegend, San Diego, CA, USA), AlexaFluor700-anti-CD27, FITC-anti-IgM, PE-anti-CD21 (BD-Biosciences), PerCP-eFluor710-anti-CD1c (eBioscience, San Diego, CA, USA). Sorted live CD19+CD1c+IgM+CD27+CD21hiCD10-mature MZ and CD19+CD1c+IgM+CD27+CD21loCD10+ MZp B-cells were >95% pure. Total RNA was extracted using RNeasy Micro Kit (Qiagen) according to the manufacturer’s instructions. RNA integrity was validated using a RNA Pico Chip on the Agilent BioAnalyzer 2100, and RNA was sent to the IRIC Genomics Core Facility for RNA-seq transcriptomic profiling and analysis. Libraries were prepared using Clontech Ultra Low RNA SMARTer v4 (Takara) and sequenced on a HiSeq2000. Genes with false discovery rate (FDR) values <0.05 were considered to be differentially expressed. Gene expression levels were compared using raw read counts and the negative binomial distribution model implemented in DESEq2 [26], a differential expression analysis package developed for R, which adjusts for sample variations with the assumption that the vast majority of genes should have correlating expression levels. More specifically, the regularized log transformation (rlog) implemented in DESeq2 was used to transform raw data into log2 (readcount) for analysis and visualization.

### Volcano plot and Gene Set Enrichment Analyses (GSEAs)

The volcano plot was created using Python 3.9.5. First, a comma separated value (CSV) file containing the raw RNA-Seq gene expression data was imported into a pandas 1.3.1 DataFrame. Each gene was assigned the colour blue if significantly upregulated/downregulated (p <0.05 and fold change > or < 1), otherwise it was assigned the colour grey. Annotations for the genes were assigned black if significantly upregulated/downregulated beyond the specified p-value and n-fold change. Otherwise, they were assigned red labels regardless of their p-values or n-fold-changes if they appear in a specified list of genes to highlight. Finally, the volcano plot was generated using Matplotlib 3.4.2. Specified p-values and n-fold lines were drawn as blue lines. The total number of upregulated/downregulated genes were displayed in the upper right/left corners, respectively.

Gene Set Enrichment Analyses (GSEAs) were produced using the software application GSEA 4.1.0, developed by the Broad Institute, as previously published [27, 28]. Required input files are the expression data, the phenotype labels, and the gene set. The expression data file consists of a table whose first column is gene name, followed by gene description, followed by the genes’ expression values for each sample. The phenotype labels file associates a phenotype label (e.g., HIV+ or HIV-) with each sample in the group. Gene sets for this manuscript were downloaded from the Broad Institute’s Molecular Signatures Database (MSigDB) v.7.4.; they are KEGG’s mTOR pathway (KEGG_MTOR_SIGNALING_PATHWAY) and Transcription Factors’ CREB/cJUN pathway (CREBP1CJUN_01) The software generates an analysis for each gene set containing the Net Enrichment Score (NES), false discovery ratio (q-value), and p-value. Gene set is considered significantly modulated if q-value and p-value <0.05.

### Multicolor Flow-Cytometry

PBMCs from 15 HIV-uninfected controls and 10 HIV-infected progressors (without and with ART) were processed for flow-cytometry as previously described [14]. Briefly, live/dead exclusion was performed using Aqua-LIVE/DEAD Fixable Stain (Invitrogen Life technologies, Eugene, OR, USA). Non-specific binding sites were blocked using FACS buffer (1x PBS, 2% hi-FBS, and 0.1% sodium azide) supplemented with 20% hi-FBS, 10 μg mouse IgG (Sigma-Aldrich, St-Louis, MO, USA) and 5 μg Human BD FCBlock (BD Biosciences). The following mouse anti-human conjugated mAbs were used to detect extracellular markers on B-cells: APC Anti-CD19, BB515 Anti-IgM, BV421 Anti-CD10, BUV395 Anti-CD73, BV786 Anti-CD39, PE-Cy7 Anti-CD83 (BD Biosciences), PerCP-eFluor 710 Anti-CD1c (eBioscience, San Diego, CA, USA). For means of verifying MZ and MZp B-cell populations as to CD21 expression levels, we had three different staining cocktails, each identical except for having a variation for the PE slot, ie 1) PE-anti-NR4A1, 2) PE-anti-NR4A3, and 3) PE-anti-CD21 to verify that the MZp were indeed CD21lo when compared to MZ. For detection of membrane BAFF expression levels, the mouse anti-human PE Anti-BAFF (clone 1D6, Invitrogen) was used. For assays with Monocyte Derived-Dendritic Cells (MoDC) (see below), the following conjugated mouse anti-human mAbs were used: PE-Cy5.5 Anti-CD11c (Invitrogen), PE-Cy7 Anti-HLA-DR, BV786 Anti-CD14 (BD Biosciences) Intra-nuclear labelling was performed using the FoxP3/Transcription Factor Staining Buffer Set (eBioscience). Non-specific binding sites were blocked using 20% hi-FBS. mAbs used were the PE-conjugated human REA clone anti-mouse NR4A1, which cross-reactivity with human has been previously assessed [14], and compared to the PE-conjugated human REA isotype control (Miltenyi Biotech), as well as the PE-conjugated mouse anti-human NR4A3 (Santa Cruz Biotechnology). Intra-cellular labelling for detection of IL-10 was performed using the Intracellular Fixation & Permeabilization Buffer Set (eBioscience) and the PE mouse anti-human Anti-IL-10 mAb (Biolegend). Cells were kept at 4°C in 1.25% paraformaldehyde until analysis. Data acquisition was performed with FACSFortessa (BD-Biosciences) for blood PBMC samples, and LSRIIB (BD-Biosciences) for tonsillar samples (described below). Analyses were done with FlowJo 10 software and GraphPad Prism. All stainings were compared to that of fluorescence minus one (FMO) values and isotype controls (see gating strategy in Supplementary Figure 1). Anti-mouse Ig(κ) Compbeads and CS&T Beads were used to optimize fluorescence compensation settings and calibrate the LSRIIB, respectively.

### Human Tonsillar B-Cells

Human tonsils from HIV-uninfected individuals, who had undergone surgical tonsillectomy, were mechanically processed and cells were cryopreserved in liquid nitrogen until use, as described above. Cells were thawed, washed in IMDM and B-cells were negatively enriched >95% by an immunomagnetic based technology (Dynabeads Untouched Invitrogen Life technologies). Total B-cells were subsequently cultured at a concentration of 10^6^ cells/mL in IMDM supplemented with 10^−4^ β-2-mercaptoethanol, 10% hi-FBS and 1% penicillin/streptomycin, in absence or presence of stimuli (PMA/ionomycin or human recombinant soluble BAFF/BLyS/TNFSF13B Protein (R&D System)) for 18 h at 37°C and 5% CO2. Cells were cultured with Brefeldin A for the last 4 hours of incubation prior to staining for intra-cellular IL-10 expression levels. Cells were recovered and processed for flow-cytometry as stated above.

### qPCR characterisation of NR4A1, NR4A3 and CD83 mRNA expression levels by Blood MZp B-cells

MZp B-cells sorted from the blood of HIV-uninfected controls, and cultured with or without soluble BAFF at 50 ng/mL or 500 ng/mL overnight, were assessed for their expression levels of NR4A1, NR4A3 and CD83 mRNA by qPCR. Briefly, blood MZp B-cells were cell-sorted as stated above and lysed with TRIzol solution (ThermoFisher). mRNA was extracted with the phenol/chloroform technique. Quality of extracted mRNA (260/280 and 260/230 ratios) was assessed with a DS-11 FX Spectrophotometer (DeNovix, Delaware, US). RNA was considered pure when 260/280 ratio was above 1.8 and 260/2360 was above 2.0. One-step qPCR was done with the QuantiNova SYBR Green qPCR Kit (QIAGEN) by using 20 ng RNA from each donor. GADPH was used as a housekeeping gene. Primers used were: NR4A1 forward: 5’-CCAGCACTGCCAAACTGGACT A-3’; NR4A1 reverse: 5’-CTCAGCAAAGCCAGGGATCTTC-3’; NR4A3 forward: 5’-CCCTTTCAGACTATCTGTACGGAC-3’; NR4A3 reverse: 5’-CTCAGTGTTGGAATGGTAAAAGAAG-3’; CD83 forward: 5’-TCCTGAGCTGCGCCTACAG-3’; CD83 reverse: 5’-GCAGGGCAAGTCCACATCTT-3’; GADPH forward: 5’ - GTCTCCTCTGACTTCAACAGCG – 3’; GADPH reverse: 5’-ACCACCCTGTTGCTGTAGCCAA – 3’; Ct values were determined with the thermocycler Corbett Research RotorGene 6000. Ct is the value of the numbers of cycles necessary for the number of RNA copies to reach a precise fluorescence threshold. The relative expression of mRNAs was determined by the ΔCt method, where Ct values of each gene are subtracted from the Ct value of the GADPH house keeping gene. mRNA expression = (2-ΔCt) x 10000.

### Breg Functional Assays

#### For assays with tonsillar populations

As previously described [14], live autologous MZp B-cells, CD1c^-^ B-cells (negative control) and CD4^+^ T-cells were sorted (see description above for RNA-Seq) from human tonsillar samples of HIV-uninfected donors. CD4^+^ T-cells were cultured alone or co-cultured with either of the B-cell populations at a ratio of 3:1 for 36 h at 37°C and 5% CO2, on anti-CD3 (2 μg/mL) (ULTRA-LEAF Biolegend) coated flat bottomed 96 well plates with soluble anti-CD28 (2 μg/mL) (ULTRA-LEAF Biolegend), and with or without human recombinant soluble BAFF at 500 ng/mL. The read-out of Breg function was based on cell cycle entry of CD4^+^ T-cells, which was assessed by flow-cytometry analyses of intra-nuclear Ki67 expression levels using the PE-mouse anti-human Ki67 mAb (eBioscience), as described above.

#### For assays with blood cell populations

Because of limited quantities of cells in our populations of interest, we proceeded to High Content Screening (HCS) assays, which read-out for Breg function was based on mortality of CD4^+^ T-cells rather than cell cycle entry. As described above, autologous MZp B-cells, CD1c^-^ B-cells (negative control) and CD4^+^ T-cells were sorted (see description above for RNA-Seq) from human PBMCs of HIV-uninfected controls and HIV-infected progressors (without and with ART). CD4^+^ T-cells were cultured alone or co-cultured with either of the B-cell populations at a ratio of 3:1 for 60 h at 37°C and 5% CO2 on anti-CD3 (2 μg/mL) (ULTRA-LEAF Biolegend) coated flat bottomed 384 CellCarrier Ultra microplates (PerkinElmer) with soluble anti-CD28 (2 μg/mL) (ULTRA-LEAF Biolegend) in the presence or absence of a mouse anti-human PD1 (2 μg/mL) blocking antibody (Biolegend). CD4^+^ T-cells were identified using a mouse-anti human CD4 (OKT4 clone) primary mAb and FITC goat anti-mouse secondary antibody (Invitrogen). The cells were then fixed using 4% paraformaldehyde before staining the nuclei with Hoescht 33342 (Invitrogen). Data acquisition was performed with Operetta and analyses done using Harmony and GraphPad Prism. The algorithm excluded debris based on Hoescht intensity, then excluded B-cells based on FITC negativity and dead cells were counted based on the intensity of Live or Dye CellMask signals of a mortality control. To this end, CD4^+^ T-cells were cultured alone in a flat bottomed 96 well plate for 60 h in the same conditions and then collected and placed at 56°C in a water bath for 45 min. The cells were then stained with Live or Dye CellsMask for Live/Dead exclusion.

#### MoDC:B-cell co-cultures

To assess the impact of membrane-bound BAFF on MZp expression levels of Breg markers, MoDC were generated from blood samples of HIV-uninfected controls, as previously described [6]. Briefly, cryopreserved PBMCs were thawed, washed in IMDM and CD14^+^ monocytes were negatively enriched >95% by an immunomagnetic based technology (Dynabeads Untouched Invitrogen Life technologies). Total CD14^+^ monocytes were subsequently cultured at a concentration of 5×10^5^ cells/mL in IMDM supplemented with 10^-4^ β-2-mercaptoethanol, 10% hi-FBS, 1% penicillin/streptomycin, 100 ng/mL Human Recombinant IL-4 and 250 ng/mL Human Recombinant GM-CSF (ThermoFisher) for six days at 37°C and 5% CO2. During the third day of incubation, 30% of growth conditions were reinstated. At day 6, MoDC purity was assessed by flow-cytometry as stated above (>90% of cells were HLA-DR^+^CD11c^+^CD14^+/-^). MoDC were subsequently cultured, with and without LPS (2 μg/ml), overnight at 37°C and 5% CO2. MoDC were then assessed for membrane BAFF expression levels by flow-cytometry as stated above, fixed in 0,25% paraformaldehyde, washed and co-cultured with autologous total B-cells (enriched as described above) at a 1:10 ratio overnight at 37°C and 5% CO2. Lastly, B-cells were harvested and processed for flow cytometry as stated above.

### Statistical Analyses

Statistical significance of differences between groups was assessed with a one-way ANOVA with post-hoc Tukey test for data normally distributed or otherwise with Kruskal-Wallis test with post-hoc Dunn test. Statistical significance of difference in clinical data between HIV-infected progressors before and after ART was assessed with the Wilcoxon matched-pairs test. Statistical significance of difference between HIV-infected progressors and EC was assessed with the Mann-Whitney U test. Analyses were performed using GraphPad Prism 9.1.1, on Windows. Results were considered significant when p < 0.05.

## Results

### Socio-demographic and data characteristics of the cohorts used in this study

Characterizations of the samples from the PHI cohort selected for this study show that there are significant differences for plasma viral loads and blood CD4^+^ T-cell counts between HIV-infected progressors before and after ART treatment, as expected. No significant difference for age, CD4^+^ T-cell counts and CD8^+^ T-cell counts were found between the HIV-infected progressors (5-8 months) and EC (Table 1).

### Blood MZp B-cells from HIV-infected progressors show an altered transcriptomic profile

We have previously shown that HIV-infected progressors from the Montreal PHI cohort presented higher levels of both BAFF and MZp B-cell frequencies in their blood, despite 1 year of ART [3]. To further assess the impact of HIV infection and associated excessive BAFF condition on deregulation of MZp, we FACS-sorted MZ and MZp B-cell populations from the blood of 3 PHI HIV-infected progressors (5-8 months), 3 ECs and 3 HIV-uninfected controls selected from the above mentioned study [3], and proceeded to transcriptomic analyses by RNA-seq. As depicted in Figure 1, PCA analyses of the top 5000 varying genes show that the gene expression profiles of blood MZ and MZp B-cell populations from the HIV-infected progressors present a distinct pattern when compared to that of both EC and HIV-uninfected controls who share greater similarity (Fig. 1A). Given this similarity between EC and HIV-uninfected controls and the scope of this manuscript, we then generated a heat map of the top 100 differently expressed genes (DEG) between blood MZp B-cells of the HIV-infected progressors and HIV-uninfected controls. We show that among those, Interferon Stimulated Genes (ISG) such as OAS1 (p = 3×10^-6^), SAMHD1 (p = 5×10^-6^) and SP110 (p = 3×10^-10^) are significantly upregulated in MZp from the HIV-infected progressors, whereas genes that have a role in regulation are severely downregulated in MZp from these same individuals (Fig. 1B and Fig. 2). The DEG were also analysed by a volcano plot, showing that 709 genes are significantly downregulated and 392 are significantly upregulated in blood MZp B-cells from the HIV-infected progressors (Fig. 1C) when compared to HIV-uninfected controls. Gene Set Enrichment Analyses (GSEA) show that both PI3K-AKT-mTOR and CREB/cJUN signatures are downregulated in blood MZp B-cells from the HIV-infected progressors (Fig. 1D and 1E, and Supplementary Figure 2). Accordingly, blood MZp B-cells from the HIV-infected progressors express higher levels of gene transcripts for exhaustion markers such as T-bet (p = 0,1) and CD11c (p = 0,009), FCRL5 (p = 0,003) and CD85j (p = 0,007), as well as the negative regulators CD22 (p = 0,05) and CD72 (p = 0,003) (Fig. 1F-K) when compared to blood MZp B-cells from HIV-uninfected controls. Altogether, these results suggest that HIV disease progression greatly solicits the activities of MZ B-cell populations, which leads to changes in their transcriptomic profile and possibly functional impairment, especially in the MZp B-cell sub-population, which present signs of exhaustion.

**Figure 1.**
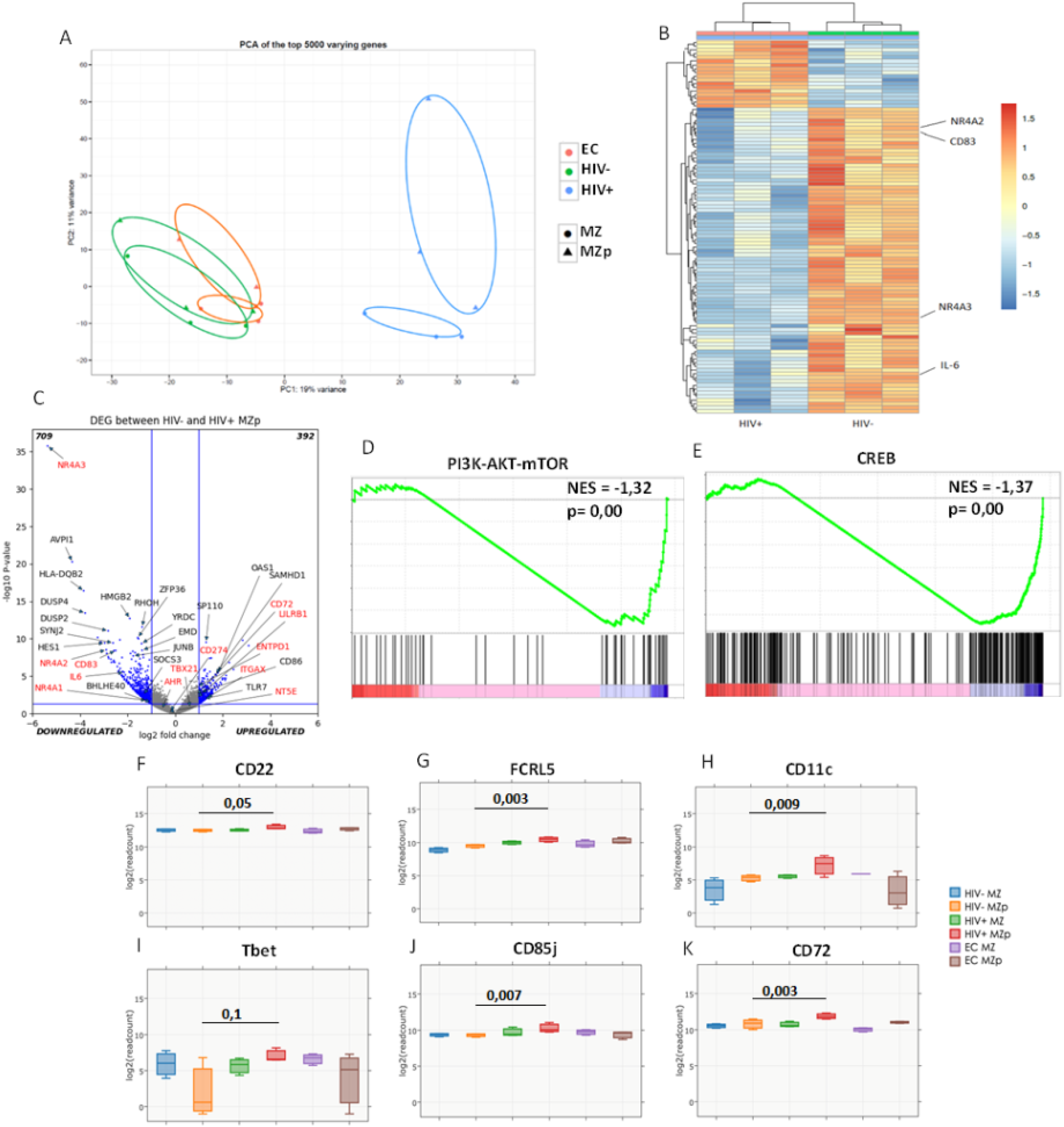
The transcriptomic profile of blood marginal zone (MZ) B-cell populations from HIV-infected progressors is different from that of elite controllers and HIV-uninfected controls. Data exploration of the transcriptomic analyses by RNAseq of sorted mature marginal zone (MZ) and MZ precursor-like (MZp) B-cells from the blood of 5-8 months HIV-infected progressors (HIV+), elite controllers (EC) and HIV-uninfected controls (HIV-). Principal Component Analysis (PCA) of the top 5000 genes (**A**). Heat Map of the top 100 Differentially Expressed Genes (DEG) between MZp B-cells from HIV-infected progressors and HIV-uninfected controls (full-sized Heat Map in Supp. Materials) (**B**). Volcano Plot between blood MZp B-cells from the HIV-uninfected and HIV-infected progressors groups (**C**). Gene Set Enrichment Analyses (GSEA) of PI3K-AKT-mTOR (NES = −1,32, p < 0,0001) (**D**) and CREB/cJUN (NES = −1,37, p < 0,0001) (**E**) pathways in blood MZp B-cells from HIV-infected progressors when compared to HIV-uninfected controls. RNAseq analyses of Tbet (**F**), CD11c (**G**), FCRL5 (**H**), CD22 (**I**), CD72 (**J**) and CD85j (**K**) gene expression levels. N = 3 for each group of participants. The Wald Test with Benjamini-Hochberg correction was used for RNAseq analysis (Fig. 1F-K)

**Figure 2.**
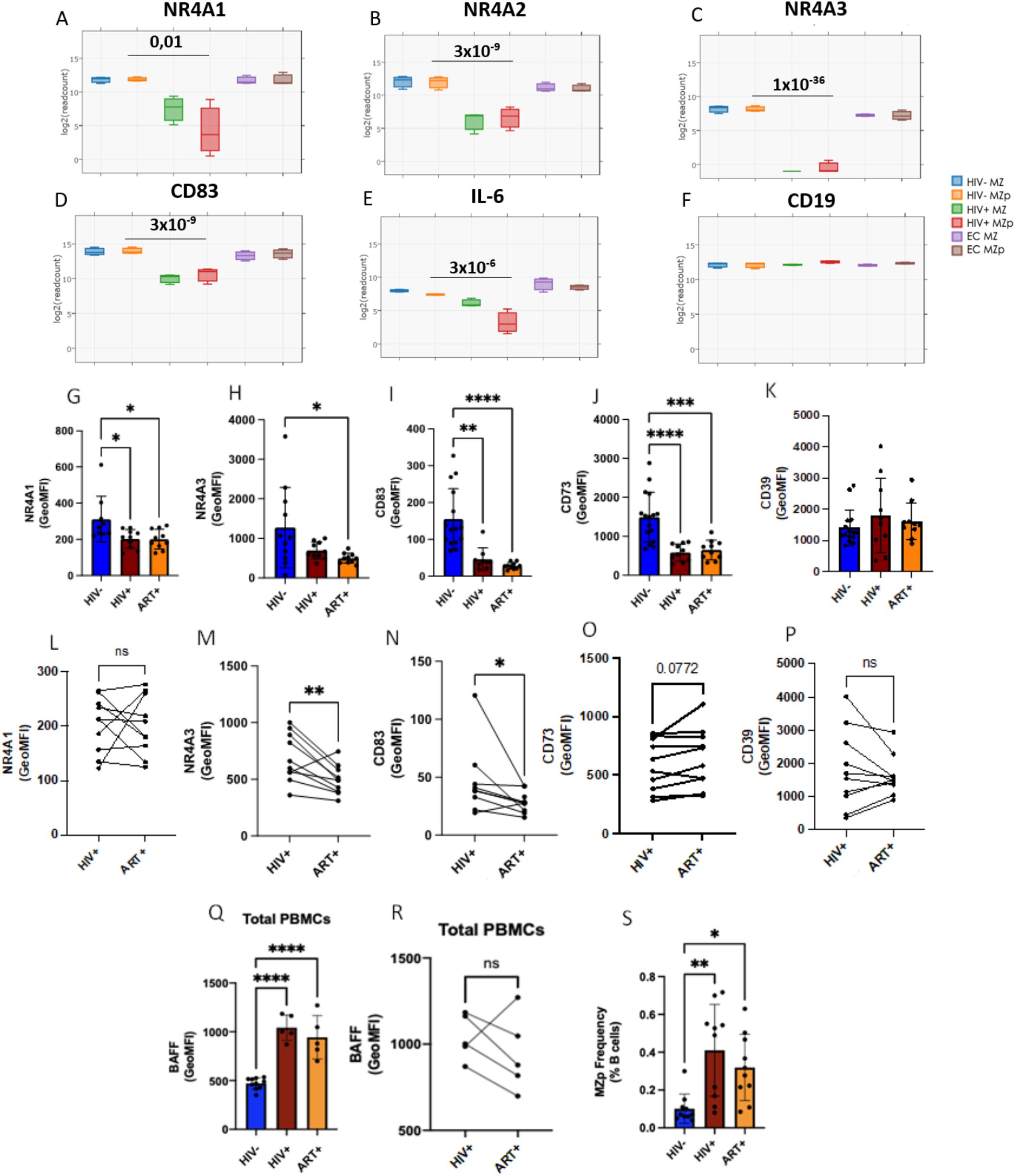
The expression levels of the Breg markers NR4A1, NR4A2 and NR4A3 are significantly downregulated in precursor-like marginal zone (MZp) B-cells from the blood of HIV-infected progressors. RNAseq analyses of MZ precursor-like (MZp) from the blood of HIV-uninfected controls (HIV-), 5-8 months HIV-infected progressors (HIV+) and elite controllers (EC) demonstrating gene expression levels of NR4A1 (**A**), NR4A2 (**B**), NR4A3 (**C**), CD83 (**D**), IL-6 (**E**) and CD19 (**E**), the latter which was used as a baseline strongly expressed B-cell gene (n=3 for each study group). Multicolour Flow-Cytometry analyses of protein expression levels of the Breg markers NR4A1, NR4A3, CD83, CD39 and CD73 were assessed on blood MZp B-cells, gated from total PBMCs of 15 HIV-uninfected controls and 10 HIV-infected progressors before and after ART. Shown are NR4A1 (**G**), NR4A3 (**H**), CD83 (**I**), CD73 (**J**) and CD39 (**K**) expression levels. Comparison between NR4A1 (**L**), NR4A3 (**M**), CD83 (**N**), CD39 (**O**) and CD73 (**P**) expression levels between primary infection and following ART for the same donors. Membrane BAFF expression levels were assessed on total PBMCs of 10 HIV-uninfected controls and 5 HIV-infected progressors before and after ART (**Q**). Comparison between membrane BAFF expression levels on total PBMCs of HIV-infected progressors before and after ART for the same donors (**R**). Relative frequencies of MZp in the blood of 10 HIV-uninfected controls and 10 HIV-infected progressors before and after ART (**S**). Protein expression levels were assessed by Geometric Mean of Fluorescence Intensity (GeoMFI). *P < 0,05; ** P< 0,01; *** P< 0,001; ****P< 0,0001. The Wald Test with Benjamini-Hochberg correction was used for RNAseq analysis (Fig. 2A-F). Normality was assessed with the Shapiro-Wilk test. A One-Way ANOVA with a Tukey post-hoc test was used for testing statistical differences between groups in Fig. 2I, J. A Kruskal-Wallis nonparametric test with Dunn’s post-hoc test was used to assess statistical differences between groups in Fig. 2 G, H, K, Q and S. A paired T-test was used in assessing differences between HIV-infected individuals before and after ART in Fig. 2.L-P and R.

### Blood MZp B-cells from HIV-infected progressors express lower levels of regulatory markers

Consistent with the severely altered profiles described above, we found highly significant downregulation of gene transcripts for the NR4A family members in blood MZp B-cells of the HIV-infected progressors. Indeed, gene expression levels of NR4A1 (p = 0,01), NR4A2 (p = 3×10^-9^), NR4A3 (p = 1×10^-36^) as well as genes that they regulate directly such as CD83 (p = 3×10^-9^) and IL-6 (3×10^-6^) (Fig. 2A-E) were very significantly downregulated in blood MZp B-cells from the HIV-infected progressors when compared to that of HIV-uninfected controls. Again, given the similarity observed with NR4As gene expression levels of HIV-uninfected controls and EC, and the scope of this manuscript, further characterization of MZp Breg capacities were conducted with HIV-infected progressors and HIV-uninfected controls. We next sought to assess whether the transcriptomic downregulation of Breg markers was suggestive of their protein downregulation. To this end, blood samples from 15 HIV-uninfected controls and 10 HIV-infected progressors (before and after ART) were selected, including samples from the individuals selected for the RNA-seq analyses. We have measured the protein expression levels of the Breg markers NR4A1, NR4A3, CD83, CD39 and CD73 by multicolor flow-cytometry on MZp B-cells from the blood of these individuals. As expected, protein expression levels of NR4A1, NR4A3, CD83 and CD73 were downregulated by blood MZp B-cells from HIV-infected progressors when compared to those of HIV-uninfected controls (Fig. 2G-K). Strikingly, ART did not restore expression levels of these Breg markers and some of them, such as NR4A3 and CD83 were further downregulated despite ART (Fig. 2L-P). Of note, consistent with that reported previously, membrane BAFF levels were found to be in excess in samples from these HIV-infected progressors and were not restored by ART (Fig. 2Q-R). Also, as expected, MZp frequencies in the blood of these HIV-infected progressors, without and with ART, were significantly higher when compared to that of HIV-uninfected controls (Fig. 2S). However, no significant correlation between MZp frequencies and BAFF levels could be observed for HIV-infected progressors (R: 0,1; p value 0,95) or after ART treatment (R: 0,3; p value 0,6833). Of note, the small numbers of samples in our groups did not allow us to reach significance for correlations between blood BAFF levels and MZp frequencies, however as stated earlier, we have previously reported excess BAFF to be concomitant with increased MZp frequencies in several cohorts and study systems. Our data suggest that MZp Breg capacities may be altered in HIV-infected progressors, regardless of therapy. These data also suggest that excess BAFF might contribute to altering the Breg profile of MZp.

### The Breg function of blood MZp B-cells from HIV-infected progressors is altered, despite ART

Given the important downregulation of Breg markers by blood MZp B-cells from HIV-infected progressors, and despite ART, we next sought to investigate whether the Breg function of MZp B-cells is altered in these individuals. In order to manage with the low cellular yields of MZp B-cells in blood, High Content Screening (HCS) by fluorescence in a confocal mode was used to assess the Breg function of sorted MZp B-cells on activated autologous CD4^+^ T-cells (Fig. 3 A-D). We observed for HIV-uninfected controls that while cell count, as determined by the amount of positive CD4^+^ cells in each analysed well (indicator of proliferation), seemed to be the same when CD4^+^ T-cells were cultured alone or co-cultured with either of MZp or CD1c^-^ B-cells (negative control) (Fig 3E), mortality rates of CD4^+^ T-cells were strikingly higher in the MZp:T-cell co-culture, suggesting regulation by controlled cell death (Fig. 3F). As such, induction of apoptosis is a common regulatory function, and has been described for Tregs [29]. Accordingly, we found that MZp B-cells strongly express PD-L1 when compared to other B-cell subtypes (Supplementary Figure 3). Furthermore, the addition of an anti-PD-1 blocking antibody to the MZp:T-cell co-culture diminished the mortality rates of CD4^+^ T-cells, thus supporting a role for the PD-1/PD-L1 pathway in MZp Breg function (Fig. 3H). However, when assessing the Breg function of blood MZp from HIV-infected progressors, our results rather suggest that there is a lower induction of CD4^+^ T-cell mortality in presence of MZp, and strikingly, despite ART (Fig. 3G). Altogether, our data suggest that HIV-infection affects the Breg function of blood MZp B-cells, and this is not restored by ART.

**Figure 3.**
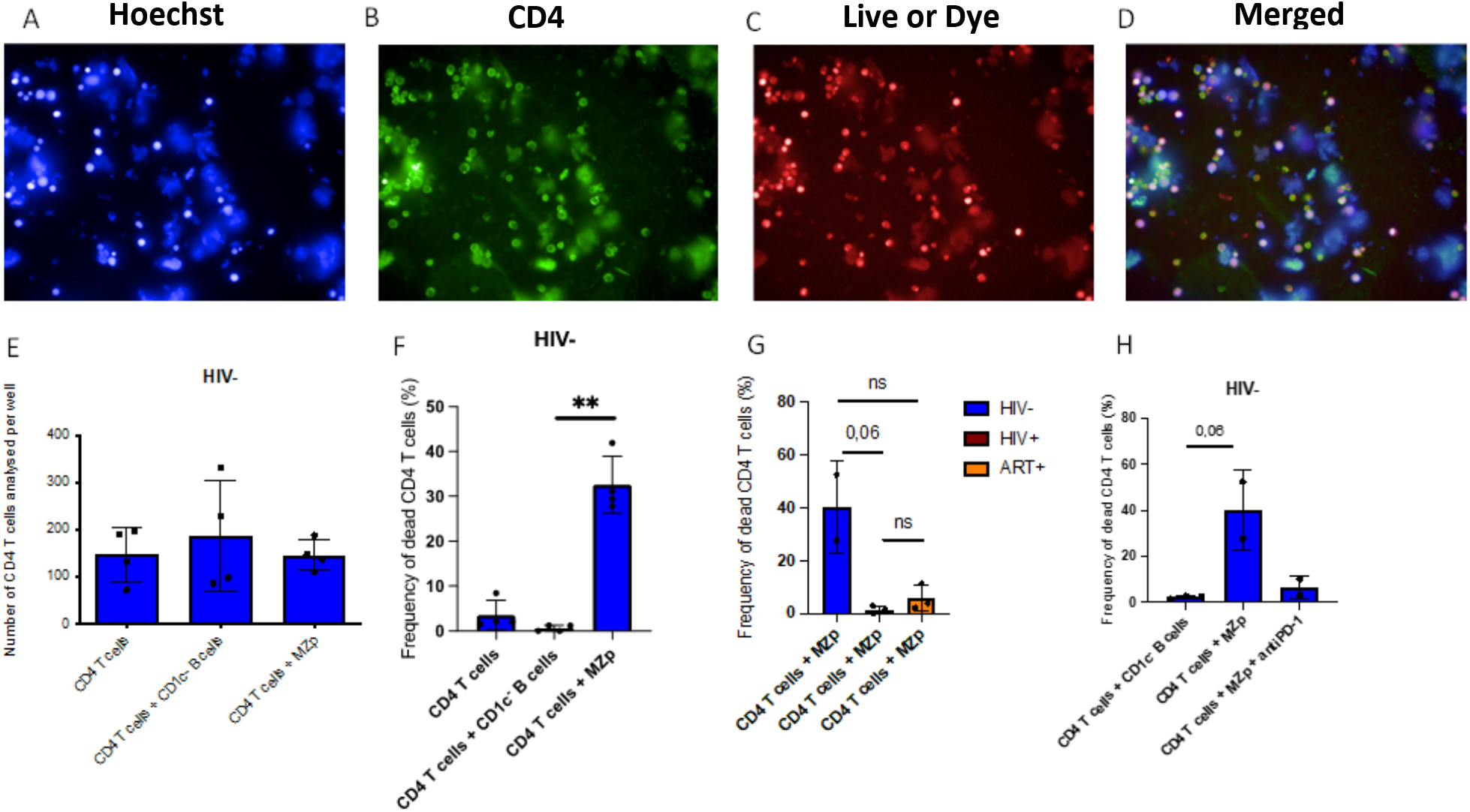
Impaired Breg function of blood precursor-like marginal zone (MZp) B-cells from HIV-infected progressors is not restored by ART. Breg function of MZp, as assessed by High Content Screening (HCS). Shown are cell staining by Hoersch (**A**), CD4^+^ T-cells staining by FITC (**B**), Live or Dye viability stain (**C**) and merged stains (**D**). CD4^+^ T-cell count on each analyzed well (**E**). CD4^+^ T-cells mortality following culture with Medium Alone or when co-cultured (3:1 ratio) with CD1c-B-cells (negative control) or MZp (**F**). CD4^+^ T-cells mortality when co-cultured (3:1 ratio) with MZp B-cells from HIV-uninfected controls and HIV-infected progressors (with and without ART) (n=3) (**G**). CD4^+^ T-cells mortality when co-cultured with MZp from HIV-uninfected controls with and without the addition of an anti-PD-1 blocking antibody at 2 μg/mL (**H**). Mortality was assessed as the relative frequency of Live or Dye cells, when compared to the total CD4^+^ T-cells. * p < 0,05; ** p< 0,01; *** p< 0,001; **** p < 0,0001. Statistical differences between groups were assessed with the Kruskal-Wallis test with Dunn’s post-hoc test. Normality was assessed with the Shapiro-Wilk test.

### Excess BAFF directly contributes to the altered Breg potential of MZp B-cells

Our RNA-seq analyses show that gene expression levels of BAFF-R are significantly down-regulated by MZp from the blood of HIV-infected progressors when compared to HIV-uninfected controls, while those of TACI and BCMA are not significantly affected (Supplementary Figure 4), albeit a trend for BCMA up-regulation by blood MZp from HIV-infected progressors could be observed. These are consistent with our observations that expression levels of BAFF-R, but not those of TACI, are decreased on B-cells following culture with soluble BAFF (Supplementary Figure 5). To assess the relative contribution of excess BAFF on deregulation of MZp Breg capacities, blood MZp B-cells from uninfected controls have been sorted and cultured in presence or absence of soluble BAFF at 50 ng/mL or 500 ng/mL overnight. Cells were then harvested for RNA extraction and a one-step qPCR for NR4A1, NR4A3 and CD83 mRNA quantification was performed, with GADPH as a housekeeping gene. Our data show that at high concentration (500 ng/mL), BAFF-treated blood MZp B-cells express lower NR4A1, NR4A3 and CD83 mRNA levels when compared to unstimulated MZp B-cells, similarly to that of sorted MZp B-cells from the blood of HIV-infected individuals (Fig. 4A-C). Likewise, we enriched total B-cells from the tonsils of HIV-uninfected donors and cultured them overnight with soluble BAFF at increasing concentrations. We found that tonsillar MZp B-cells treated with high BAFF concentrations show lower expression levels of NR4A1 and CD83 proteins when compared to those treated with lower BAFF concentrations or medium alone (Fig. 4D-G). Interestingly, we found that as soluble BAFF concentrations increase, IL-10 expression levels decrease (Fig. 4 H, I). Of note, we had previously shown that IL-10 is highly expressed by *ex vivo* unstimulated blood MZp B-cells, when compared to other B-cell populations ([24], and Supplementary Figure 6). Upon performing experiments with enriched tonsillar B-cells from HIV-uninfected donors, we observed that the total tonsillar cells from different donors expressed varying levels of total membrane BAFF (Fig 4L). This difference was mainly attributable to DC-like (HLA-DR+CD11c+CD14-CD3-CD19-) and MoDC-like (HLA-DR+CD11c+CD14+CD3-CD19-) populations, as membrane BAFF expression on T-cells was similar between the donors (Fig 4 M-O). Strikingly, tonsils expressing higher total membrane BAFF levels have reduced MZp Breg function, while tonsils expressing lower BAFF levels have a MZp Breg function comparable to that expected at a 1:3 ratio [14] (Fig. 4J-K). Interestingly, this latter Breg function is altered upon addition of high levels of soluble BAFF (Fig. 4P). To further investigate the impact of membrane BAFF levels on MZp Breg capacities, blood derived MoDC stimulated with LPS prior to paraformaldehyde fixation (Supplementary Figure 7), were co-cultured overnight with autologous blood B-cells at a 1:10 ratio. Blood MZp B-cells expression of regulatory markers were then assessed by flow-cytometry (Supplementary Figure 7). We observe a trend for the downregulation of NR4A1 and NR4A3 expression levels by these blood MZp B-cells. Although this finding will need further investigation, it is tempting to speculate that DC-like and MoDC-like populations in secondary lymphoid tissues, which are known to interact with B-cell populations such as MZ B-cells [11], may influence MZp Breg activity through membrane and/or released BAFF expression levels.

**Figure 4.**
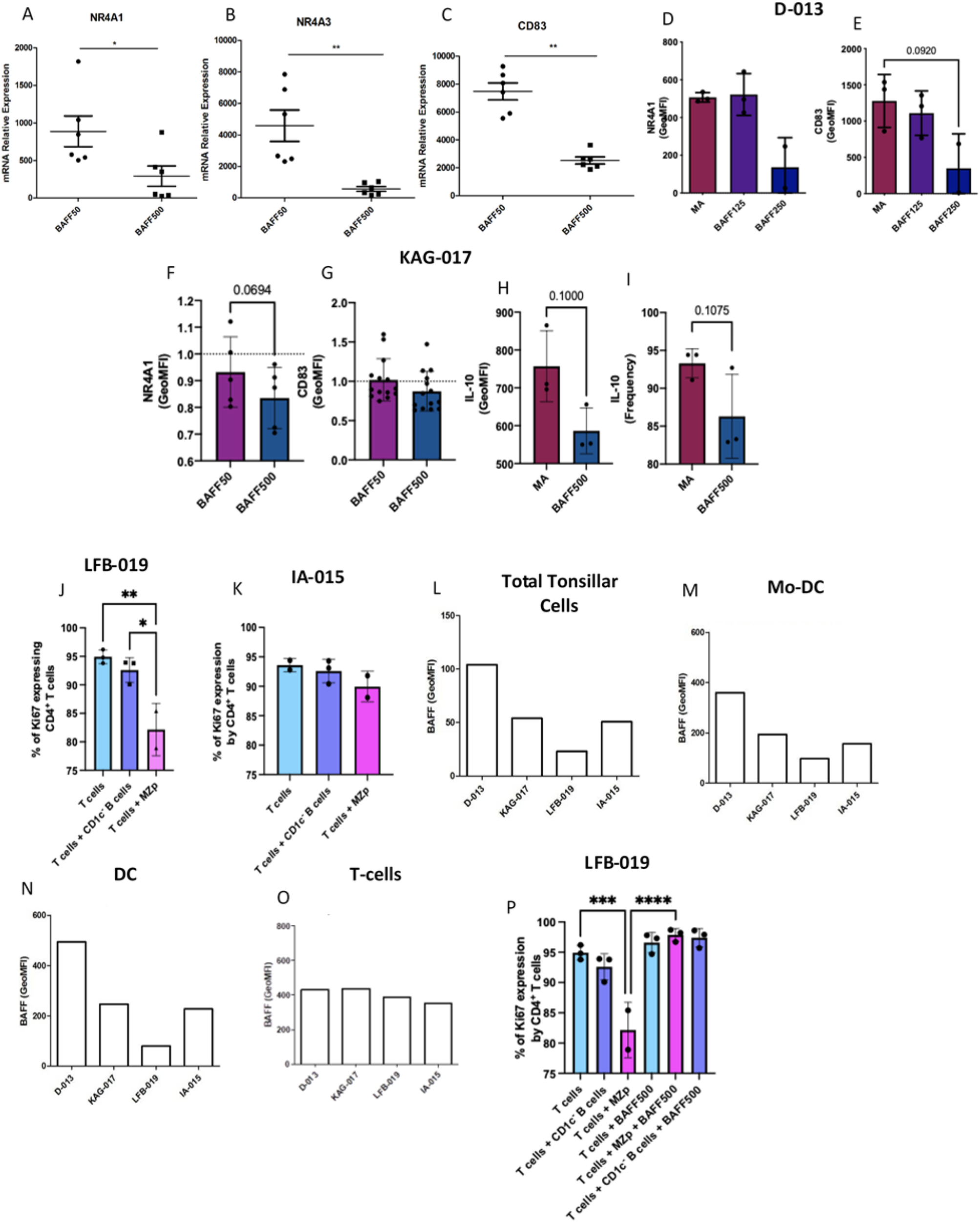
High levels of BAFF significantly downregulate precursor-like marginal zone (MZp) B-cell expression levels of immunoregulatory markers and influence their Breg function. qPCR analyses of NR4A1 (**A**), NR4A3 (**B**) and CD83 (**C**) mRNA relative expression levels by blood MZp B-cells from HIV-uninfected individuals when cultured with 500 ng/mL soluble BAFF (2 donors, n =3 each). Flow-cytometry analyses of BAFF-treated tonsillar B-cells from HIV-uninfected donor D-013 (n=3) showing NR4A1 (**D**) and CD83 (**E**) protein expression levels. Flow cytometry analyses of BAFF-treated tonsillar B-cells from HIV-uninfected donor KAG-017 (n = 5) showing NR4A1 (**F**), CD83 (**G**) and IL-10 (**H**) expression levels as well as total relative frequencies of MZp expressing IL-10 (**I**). Data of (**F**) and (**G**) are shown as fold-over Medium Alone (MA) due to variabilities between different tonsillar samples from the same donor. Breg function of MZp B-cells from two different tonsillar donors, LFB-019 (**J**) and IA-015 (**K**) have been assessed as relative frequencies of CD4^+^ T-cells expressing intracellular Ki67, as an indicative of cell cycle entry. Flow cytometry analyses of membrane BAFF levels from total tonsillar cells (**L**), Mo-DC (**M**), DC (**N**) and CD3^+^ T-cells (**O**) from different tonsillar donors. Breg function of MZp B-cells from LFB-019 donor after treatment with high soluble BAFF (500 ng/mL) for 18 hours, assessed as relative frequencies of CD4^+^ T-cells expressing intracellular Ki67 (**P**). Relative frequencies of MZp expressing IL-10 were assessed relatively to the percentage of total MZp B-cells. Relative frequencies of CD4^+^ T-cells expressing Ki67 were assessed relatively to the percentage of total CD4^+^ T-cells. Expression levels were assessed with Geometric Mean of Fluorescence Intensity (GeoMFI). *P < 0,05; ** P< 0,01; *** P< 0,001; ****P< 0,0001; MA - Medium Alone; BAFF50 - BAFF at 50 ng/mL; BAFF500 - BAFF at 500 ng/mL. Statistical differences between groups were assessed with the Kruskal-Wallis test with Dunn’s post-hoc test. Normality was assessed with the Shapiro-Wilk test.

## Discussion

We have previously shown that excess BAFF in the blood of HIV-infected progressors was concomitant with deregulation of the blood B-cell compartment, notably with increased frequencies of MZp B-cells [3], despite ART. In this study, we report that blood MZ populations, especially MZp B-cells from HIV-infected progressors present a different transcriptomic profile when compared to that of ECs and HIV-uninfected controls, suggestive of their being highly solicited and functionally impaired, with signs of exhaustion. Importantly, differences include the significant downregulation of gene transcripts for the molecules NR4A1, NR4A2, NR4A3 and CD83, which we have previously associated with the Breg potential of MZp B-cells in HIV-uninfected individuals [14]. Furthermore, the protein expression levels of these molecules as well as those of CD73, also associated with regulatory capacities [30], are downregulated in blood MZp B-cells of HIV-infected progressors, and despite ART. Even more importantly, we found that the Breg function of blood MZp B-cells is impaired in HIV-infected and ART treated individuals. Consistent with our previous observations [3], levels of BAFF were found to be in excess in the blood of HIV-infected and ART treated individuals selected for this study, when compared to HIV-uninfected controls. Importantly, *in vitro* studies show that high levels of BAFF impact on the expression of regulatory proteins by MZp B-cells from tonsils of HIV-uninfected donors and have a direct effect on their Breg function.

Our findings are suggesting that in the context of HIV, which we characterize as a persistent excessive BAFF environment, MZ populations may be highly solicited, selected, and expanded to produce antibodies at the expense of their Breg competence. As such, BAFF levels may dictate MZ B-cell activities, from Breg to antibody production, which in the context of HIV may be expedited, bypassing the Breg capacity, for tentative eradicative purposes. As to whether MZ B-cells antibody responses are beneficial or a nuisance to the host in the context of HIV remains to be established. As such, given the facts that HIV-Tg and BAFF-Tg animals presented breakage of tolerance and autoimmunity [7, 31–33], and that excess BAFF and MZ B-cell deregulations have also been reported in the context of autoimmune diseases such as Systemic Lupus Erythematosus (SLE) and Sjögren Syndrome (SS) [34], it would be sound to bear in mind that MZ B-cell antibody responses in the context of excess BAFF may be nourishing polyclonal, possibly auto-reactive, low-affinity antibody responses, whose pertinence in the battle against HIV have yet to be determined.

Notably, individuals with SLE or SS present a strong IFN signature, which promotes BAFF expression, reminiscent of what was observed in the context of chronic infections such as with HIV [35, 36]. Thus, unsurprisingly, we found that genes that are associated with pro-inflammatory outcomes and ISG are upregulated in blood MZp B-cells of HIV-infected progressors. As such, we found that transcripts for genes involved in the regulation of IFN responsiveness such as OAS1, SP110 and IRF7 are highly upregulated in MZp B-cells of HIV-infected progressors when compared to EC or HIV-uninfected controls. These data suggest that MZp B-cells are in a hyperactive state in the context of HIV, in a similar manner to that observed in autoimmune diseases such as SLE or SS.

Our GSEA analyses show that the major PI3K-AKT-mTOR pathway is downregulated in blood MZp B-cells from HIV-infected progressors. mTOR has been shown to be important for normal MZ B-cell function, as TACI signaling converge towards mTOR in order to lower MZ activation threshold [37], which is in accordance with their role as first-line innate-like cells that respond quickly to blood borne pathogens [11]. Interestingly, the transcript for the gene coding for BAFF-R is significantly downregulated by blood MZp B-cells of HIV-infected progressors while the transcript for TACI remains highly expressed (Supplementary Figure 4). Unfortunately, we did not assess BAFF-R nor TACI protein expression levels by blood MZp in these individuals, but they are also likely to be modulated as we found that excess soluble BAFF *in vitro* downregulates cell surface expression levels of BAFF-R but not those of TACI on blood MZp (Supplementary Figure 5). It is therefore possible that in HIV-infected progressors, reduced MZp expression of BAFF-R reflect ongoing exposure to increased amounts of its cognate ligand BAFF. Moreover, our data also suggest that BAFF signals may be mostly mediated via TACI during HIV infection. Importantly, the binding of BAFF to TACI allows the recruitment of TRAFs such as TRAF3 [38], which has been shown to inhibit the CREB pathway in B-cells [39], known to induce the expression of the NR4As [40, 41], and which we find is also downregulated in MZp B-cells from HIV-infected progressors. Of note, a trend of low TRAF6 gene transcripts was found in HIV-infected individuals (p = 0,1), but not TRAF3 (Supplementary Figure 4), which could mean that TRAF3 could see a higher recruitment rate due to a lower competition for TACI signals. This data suggests that the excess BAFF found during HIV infection may lead to CREB and NR4As downregulation via excess TACI signaling, and thus excess TRAF3 recruitment. Given the importance of CREB and NR4As in controlling cell activation [42], it is possible that their downregulation allows for MZp B-cells to maintain a hyperactive state despite mTOR downregulation. The fact that gene expression levels of BCMA show a trend for upregulation by blood MZp from HIV-infected progressors when compared to HIV-uninfected controls, could suggest differentiation towards an early plasmablast stage, as they also significantly up-regulated CD38 gene transcripts (Supplementary Figure 4L).

Importantly, NR4A1, NR4A2 and NR4A3 gene transcripts were downregulated in a highly significant manner in MZp B-cells from the blood of HIV-infected individuals, with p values reaching orders of magnitude of 0,01, 10^-9^ and 10^-36^, respectively. As well, other important regulatory molecules that are directly modulated by NR4As such as CD83 and IL-6 [17] were also significantly downregulated. Consistently, blood MZp B-cells from HIV-infected individuals presented significantly lower protein expression levels of NR4A1, NR4A3, and CD83, and this was not restored by ART. The importance of NR4A transcription factors in immune regulation has been highlighted by the fact that NR4A1-3 double knockout mice develop systemic autoimmune diseases [43]. Furthermore, in humans, downregulation of NR4As has been observed in the context of autoimmunity [43] and acute myeloid leukemia [44] whereby synthetic regulation of NR4A expression is currently used for eliminating cancerous cells [45]. Also, since expression of NR4As is necessary for the maintenance of FoxP3 expression by Tregs [16], there have been reports of several studies where those genes are targeted to modulate Treg responses in cancer [45]. Moreover, increasing NR4A1 expression in myeloid cells led to a diminished T-cell activation profile [20]. It is thus reasonable to think that NR4A modulation could be envisaged in view of restoring MZp Breg capacities, possibly in conjunction with therapies reducing excessive BAFF levels, already FDA approved in the treatment of SLE.

We found similar observations for the ectonucleotidase CD73, the limiting molecule of the ADO pathway [46]. Others have reported that the diminution of CD73^+^ B-cells and CD39^+^CD73^+^ B-cells and diminution of ADO correlated with a loss of CD4^+^ T-cells and disease progression during HIV infection [47], [48]. Interestingly, the HIV context is also associated with downregulated CD39 and CD73 expression levels in Tregs, despite ART [49]. In other chronic infections, such as hepatitis, a loss of CD73 and CD39 has also been observed and associated with viral load and high levels of inflammation [50]. It is also important to remember that ADO has been found to modulate NR4As expression levels via purinergic receptors such as A2_A_ [51]. Therefore, the loss of CD39 and CD73 can contribute to a reduction of extracellular ATP conversion into ADO, which could promote a pro-inflammatory environment caused by the accumulation of extracellular ATP and a decrease of NR4A expression due to a decrease in the interaction between ADO and its receptors. Since the mTOR pathway is upregulated by the binding of ADO to its receptors [52], it is possible that the lower levels of ADO production by regulatory cell populations (MZp B-cells and others) contribute to the downregulation observed for the mTOR pathway.

Interestingly, we noticed that gene transcripts for Breg markers such as IL-10 and the IL-10 regulatory transcription factor aryl hydrocarbon receptor (Ahr), recently shown to be an important Breg marker in mice [53], were not affected in blood MZp from HIV-infected progressors, whereas others such as transcription factor basic helix-loop-helix family member e40 (BHLHE40), also involved in IL-10 regulation, were significantly diminished when compared to HIV-uninfected controls (Supplementary Figure).

Consistent with their loss of several Breg markers, we show that blood MZp B-cells from HIV-infected and ART treated individuals lose their Breg function. We have previously shown that MZp B-cells from HIV-uninfected individuals can control CD4^+^ T-cell proliferation and that this involves signals delivered via CD83 [14]. Here, we show that the control of MZp B-cells on the activation of autologous CD4^+^ T-cells may also involve controlled cell-death and PD-1/PD-L1 signalling. Accordingly, the reduced Breg function observed for blood MZp from HIV-infected progressors, without and with ART, is concomitant with their lowered expression levels of PD-L1 (Supplementary Figure 3). Of note, the differences observed between PD-L1 gene transcripts, which expression levels were not affected in blood MZp from HIV-infected progressors, and PD-L1 protein expression levels, which were decreased, suggests that protein expression of PD-L1 could be affected by post-transcriptional mechanisms. Further studies are thus needed to confirm the relative involvement of all markers engaged in MZp Breg function.. Importantly, we show that excess BAFF can directly impact on MZp Breg capacities. Indeed, gene and protein expression levels of the regulatory markers NR4A1, NR4A3 and CD83 were downregulated in presence of high soluble and membrane BAFF levels, which also had an impact on IL-10 production, which could contribute further to the loss of Breg function in these cells.

When performing *in vitro* studies with tonsils from HIV-uninfected donors, we noticed that BAFF levels varied between each donor and this influenced *in vitro* MZp Breg capacities. Variations of BAFF levels between each donor appear to be especially attributable to levels expressed by myeloid cells such as DC-like and MoDC-like populations. As such, DC are an important source of BAFF, and have been shown to interact with B-cells in T-independent manners [11] and could have thus modulated the tonsillar MZ B-cell environment. Of note, in HIV-Tg mice, we have observed that myeloid DCs accumulated in the expanded MZ of these mice [54]. We have previously published that HIV infection increases BAFF levels on DC and MoDCs via Nef [6]. Likewise, elements of microbial translocation, such as LPS were also shown to increase BAFF expression by MoDCs ([6], Supplementary Figure 7). Our findings point to BAFF as being a main contributor to the loss of MZp Breg capacities during HIV infection. Similar to that observed for SLE and other rheumatic disorders, there is growing evidence that BAFF is in excess in contexts of cancer, and could also play a role in the pathogenesis of several inflammatory chronicity such as, multiple sclerosis, and HBV infection [35]. Suggesting that excess BAFF and deregulated MZp may be important indicators of deterioration of immune-competence.

We found relatively higher gene expression levels of exhaustion markers such as FCRL5 and CD85j and negative regulators such as CD22 and CD72 in blood MZp B-cells of HIV-infected progressors when compared to HIV-uninfected controls, which suggest a higher activation index. Also, MZp B-cells express both IL-21R and TLR7 (Supplementary Figure 4), which signals are important to drive B-cell differentiation towards an “age associated”-like exhausted B-cell profile. As such, blood MZp B-cells from HIV-infected progressors also possess higher gene expression levels of T-bet and CD11c, a characteristic related to age-associated B-cells, and shared by a similar heterogeneous population found to be expanded in autoimmunity, chronic infections, antibody-mediated rejection in organ transplants [55–57], and in critically ill patients with SARS-Cov-2 [58]. Interestingly, B-cells that express T-bet and CD11c have been associated with their accumulation at extra-follicular sites and production of immunoglobulins (Ig) of poor affinity [59].

To date, many groups have identified MZ-like populations with Breg capacities vs T-cells, in non-cognate fashions, *in vitro*, in murine models and in humans (Recently Reviewed by [60]). Herein, our data suggest that such Breg capacity of MZp is greatly reduced in HIV-infected progressors, and not restored by ART. And this possibly involves excessive BAFF levels reported for these individuals. As to whether MZp can exert their Breg functions through cognate interactions with CD4+ T-cells, needs further experimentation and is beyond the scope of our manuscript but deserves to be addressed. Mouse MZ B-cell populations have been shown to migrate to T-cell zones of secondary lymphoid organs (SLO), and activate CD4+ T-cells in a HEL-dependent manner [61]. This, and the fact that MZ B-cell populations have the capacity to shuttle to follicular areas [62] and furthermore that MZ B-cells were shown to possess an atheroprotective role attributed to their NR4A1 expression [22, 23], all point to the capacity of MZ populations to be recruited to outer MZ areas of SLO, in given situations. Furthermore, MZ B-cells are able to present lipidic antigens to invariant natural killer T-cells (iNKT), in the context of CD1, which confer activation via notably, the CD40-CD40L pathway [63]. All of which supporting their influence on several cell types. The capacity of MZ B-cell populations to perform antigen presentation, Breg activity or to differentiate towards Ig production may be dictated by the overall situation and environment.

## Conclusion

There is a limitation of our study in the number of participants, and we believe adding further subjects in future will strengthen the interpretation of our data. Nevertheless, the present study sheds some light in a better understanding of the important Breg potential of MZp B-cells, and its deregulation in the context of chronic inflammation involving excess BAFF, such as encountered in HIV infection. Importantly, MZp Breg potential was not restored by therapy. Our observations are essential to help develop strategies viewed at restoring MZp immune surveillance activities. To this end, existing strategies, such as dihydroergotamine (DHE) viewed to upregulate NR4As expression levels could be envisaged to reinstate MZp Breg capacities. Also, BAFF-blocking therapies, such as Belimumab (Benlysta), which is FDA approved in treating SLE, could be contemplated to try to control BAFF levels in order to lower the inflammatory burden and restore B-cell immune competence.

## Acknowledgements

We are grateful to Mario Legault for the clinical data. We are grateful to Raphaële Lambert, Jennifer Huber, and Patrick Gendron (IRIC Genomic and Bioinformatics core facilities) for RNASeq transcriptomic and data analyses, respectively. We are grateful to Guillaume Beaudoin-Bussières for the help with the BSL3 cell culture. We are also grateful to Erik Joly for his help during troubleshooting and for establishing the algorithm for HCS experiments. We are grateful to Dre Dominique Gauchat, Philippe St-Onge and Gaël Dulude for their support with the CHUM cytometry platform. We are grateful to Audray Fortin and Elise Caron for their help in establishing the HCS protocol. We are grateful to Drs Pavel Chrobak and Bertrand Allard for fruitful discussions. And lastly, we are grateful for the individuals who graciously donated the samples required for this study.

## Author Contributions

KDL, MA, JP managed all the parameters regarding the study participants. JCC prepared the samples for RNA-sequencing (IRIC). KDL and MA performed the flow-cytometry and Breg experiments. KDL performed Breg experiments by HCS. MA performed the qPCR, and MoDC experiments. MB did the first assays with soluble BAFF and tonsillar samples. KDL, MA, JP, MR analyzed the data and wrote the article. MP and MA performed the Volcano plot and GSEA statistical analyses. KDL and MA produced graphs and layouts. JP and MR designed the concept. JPR and CT manage the Montreal LTNP and PHI cohorts, and provided all the blood samples. MCQ, NB and DEK managed and provided the human tonsillar samples. All authors revised the last version of the manuscript.

