## Supplemental Figures for "Excess BAFF Alters NR4As Expression Levels and Breg Function of Human Precursor-like Marginal Zone B-cells in the Context of HIV-1 infection"

**Supplementary Figure 1. Gating strategy for multicolour flow-cytometry analyses**. After excluding doublets and dead cells, total live B-cells (CD19^+^) were positively gated. Then, CD1c^+^ B-cells were selected, followed by IgM^+^CD27^+^ double-positive B-cells. From the latter, the CD10- population was designated as marginal zone (MZ) B-cells, while the CD10^+^ population was designated as MZ precursors (MZp).

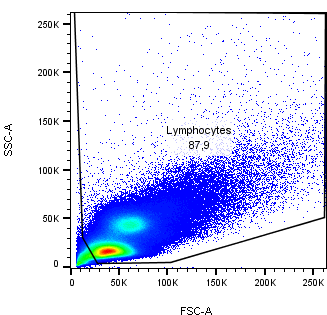

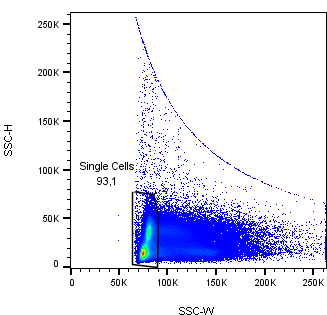

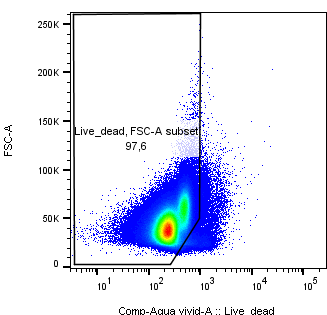

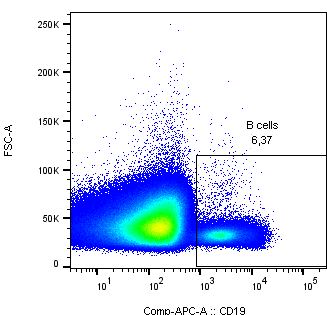

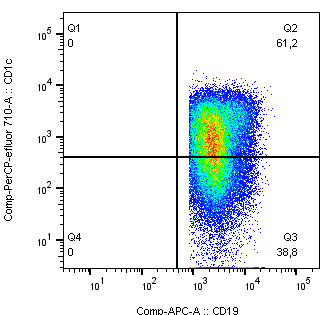

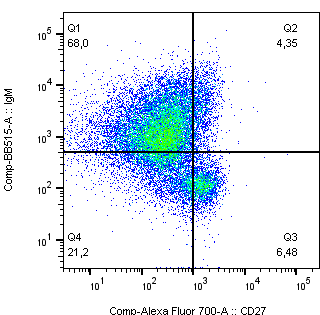

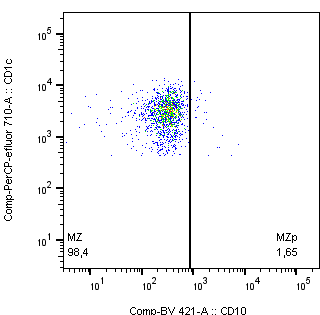

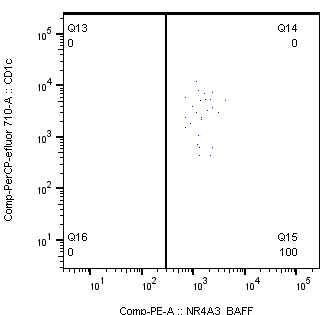

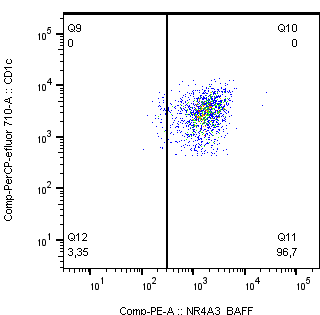

**MZp**

**MZ**

**Total B-cells**

**Supplementary Figure 2. RNAseq analyses of blood MZp B-cells PIK3-AKT-mTOR and other pathways.** Data exploration of the transcriptomic analyses by RNAseq of sorted mature marginal zone (MZ) and MZ precursor (MZp) B-cells from the blood of 5-8 months HIV-infected progressors (HIV+), elite controllers (EC) and uninfected controls (HIV-) (n=3 for each group). Shown are the gene transcripts associated with mTORC1: mTOR (**A**), MLST8 (**B**), AKT1S1 (**C**), DEPTOR (**D**), RPTOR (**E**). Gene transcripts associated with PI3K: PIK3CD (**F**), PIK3CG (**G**). Gene transcripts associated with the AKT family: AKT1 (**H**), AKT2 (**I**), AKT3 (**J**). Gene transcripts associated with the AP-1 genes: Jun (**K**), Fos (**L**). Gene transcripts associated with Protein Kinase A (PKA): PRKACA (**M**), PRKAR1A (**N**), PRKAR2A (**O**). N = 3 for each group of participants. Statistical analyses were done between blood MZp B-cells from uninfected controls (HIV- MZp) and HIV-infected progressors (HIV+ MZp). The Wald Test with Benjamini-Hochberg correction was used for RNAseq analysis.

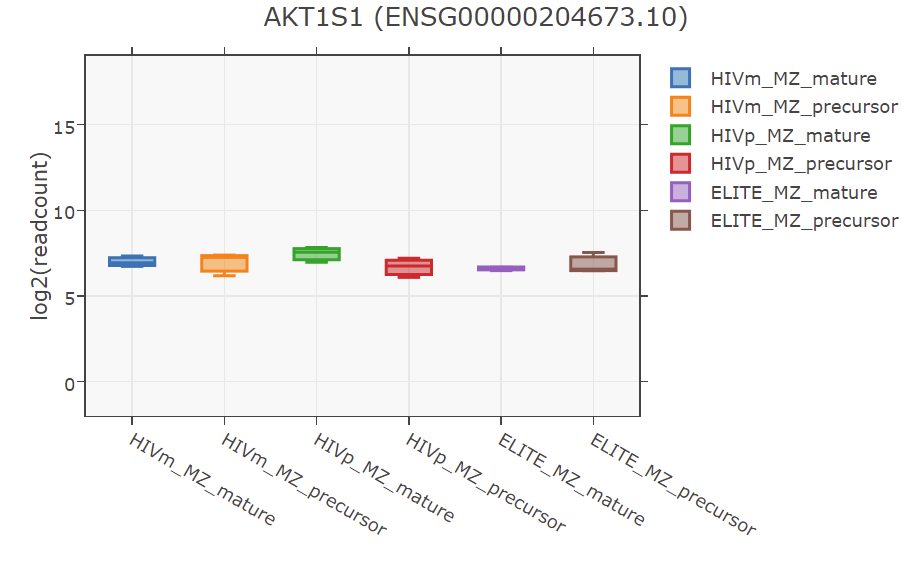

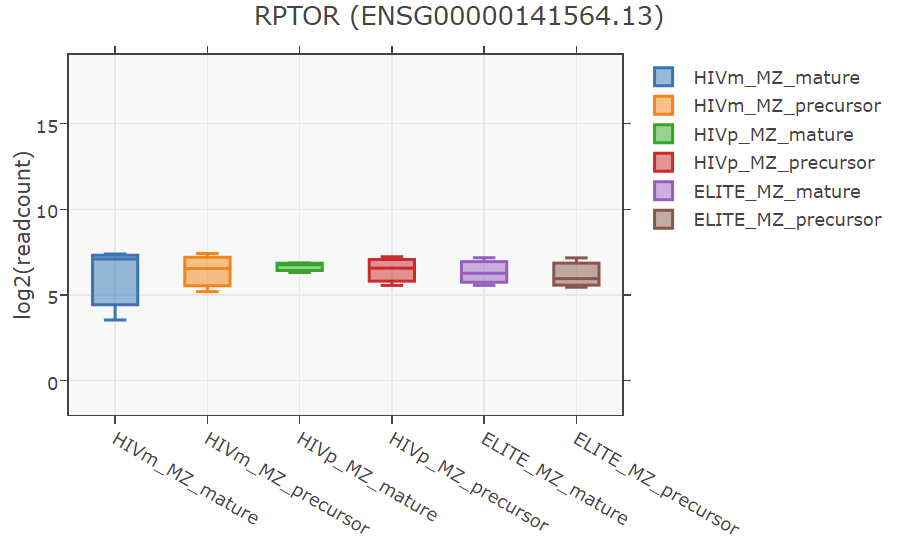

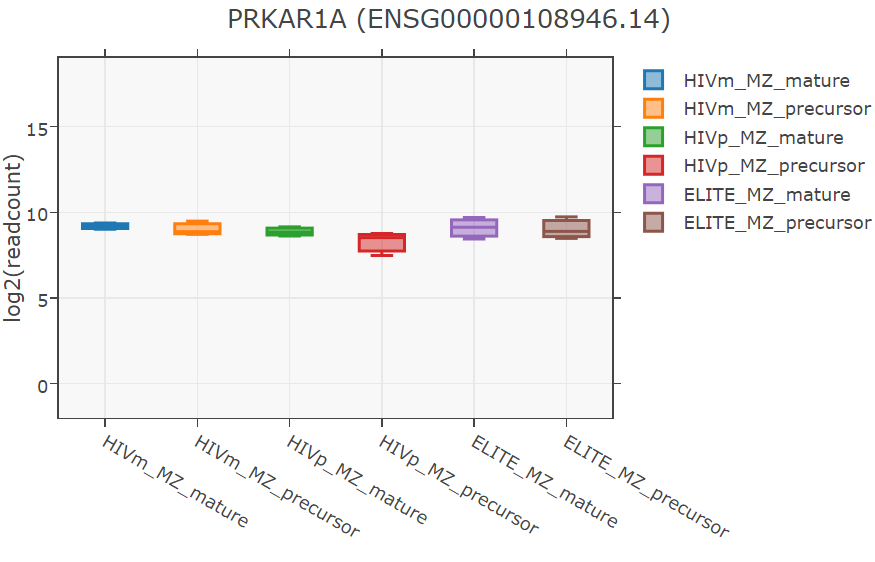

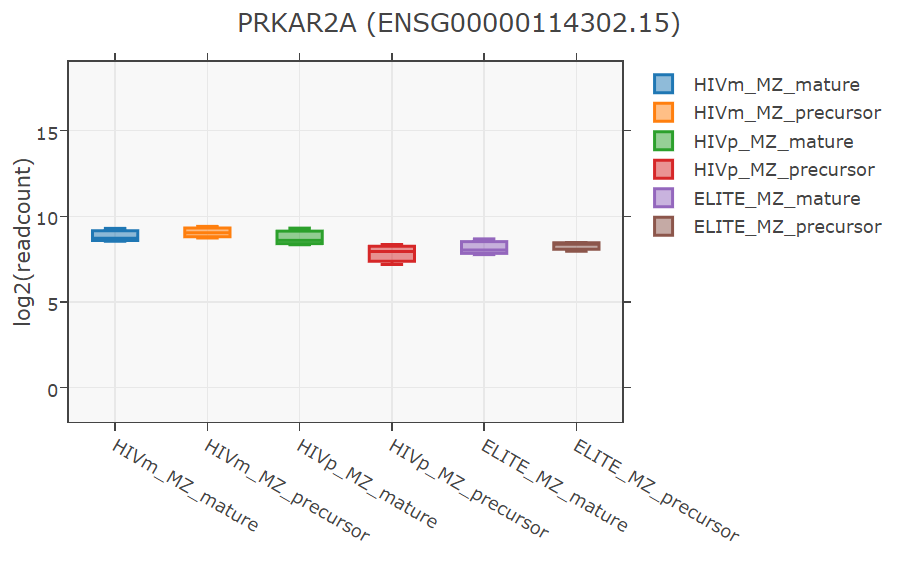

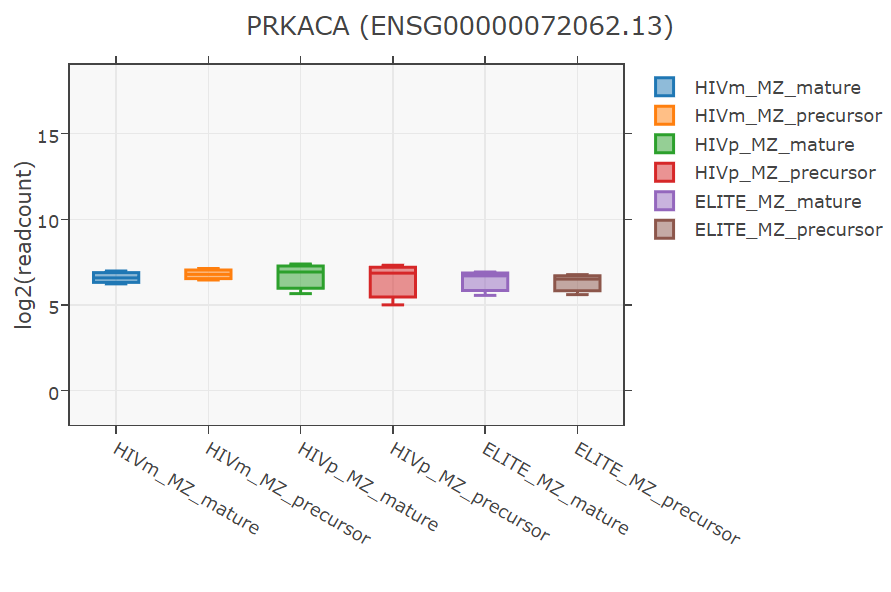

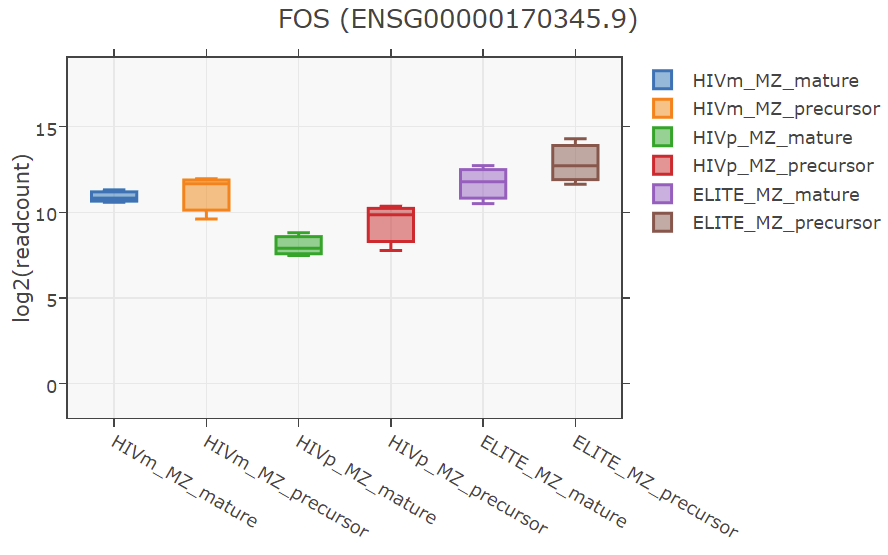

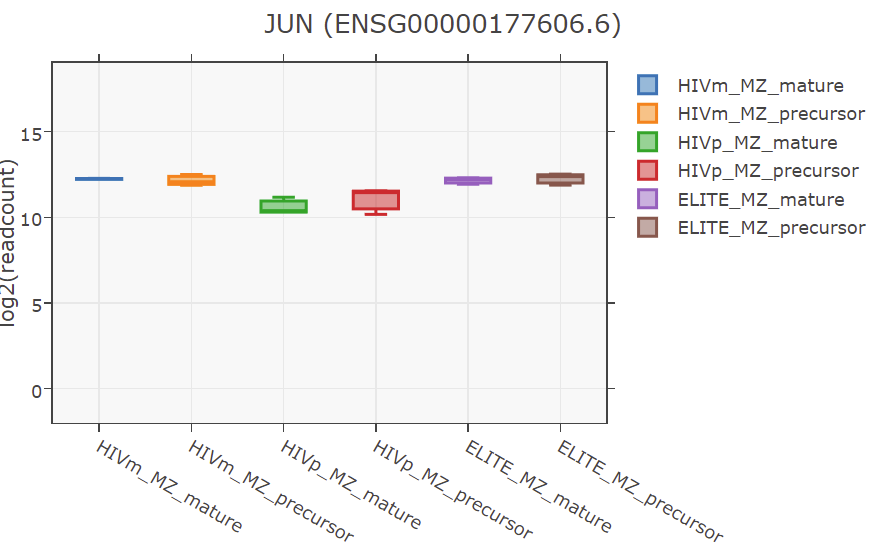

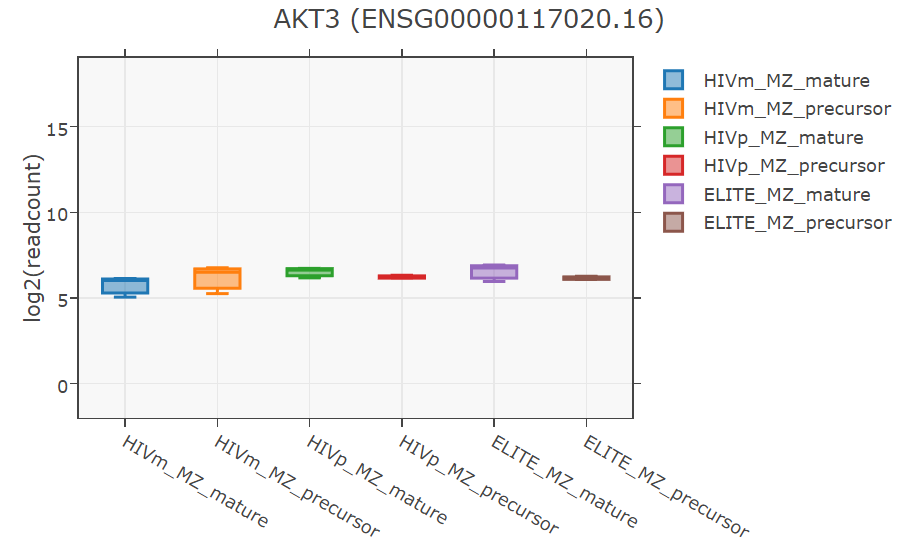

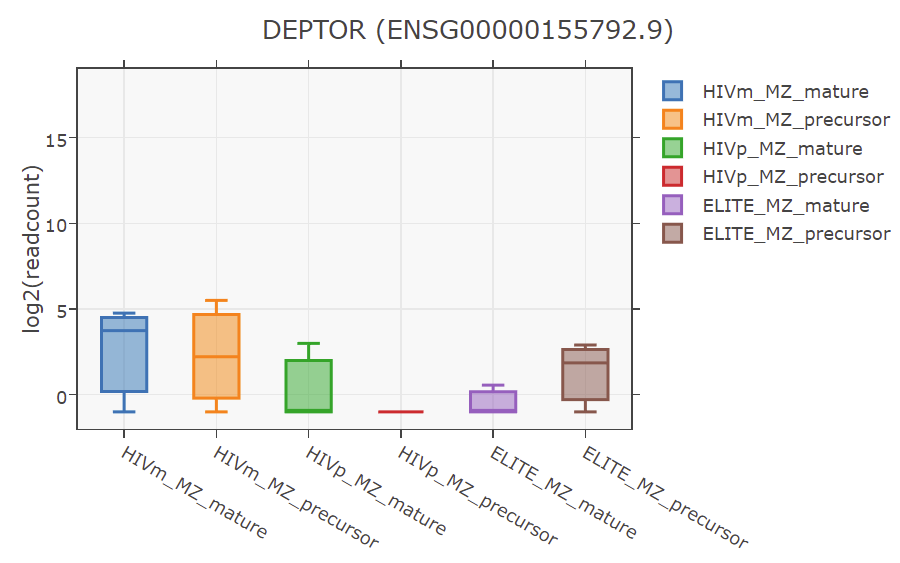

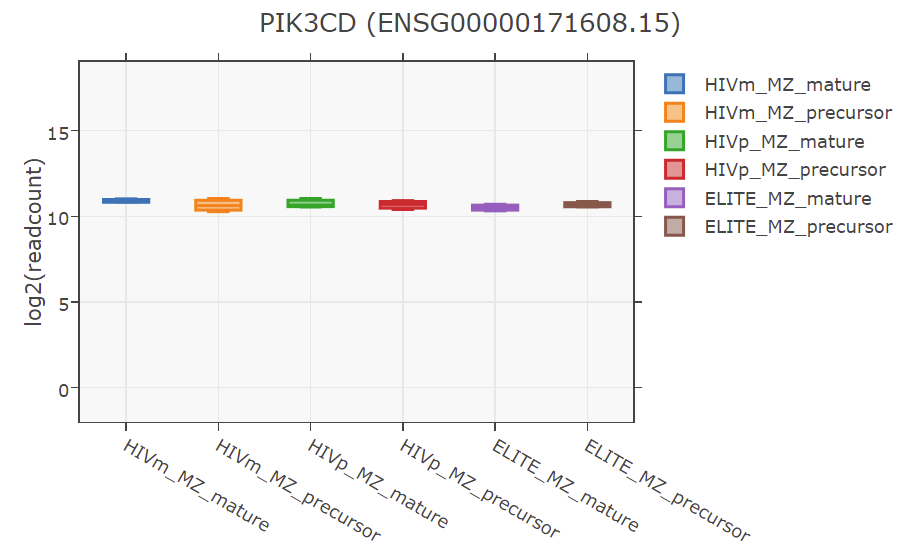

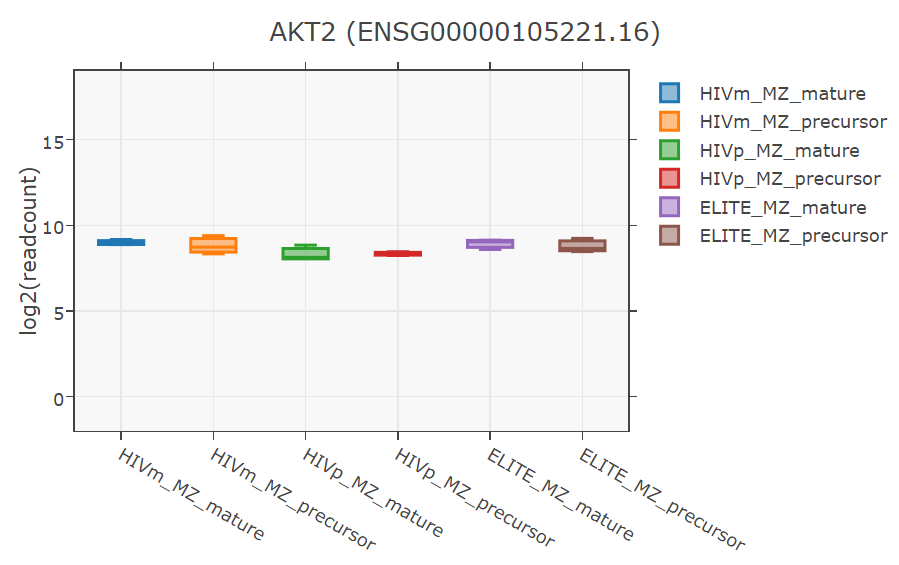

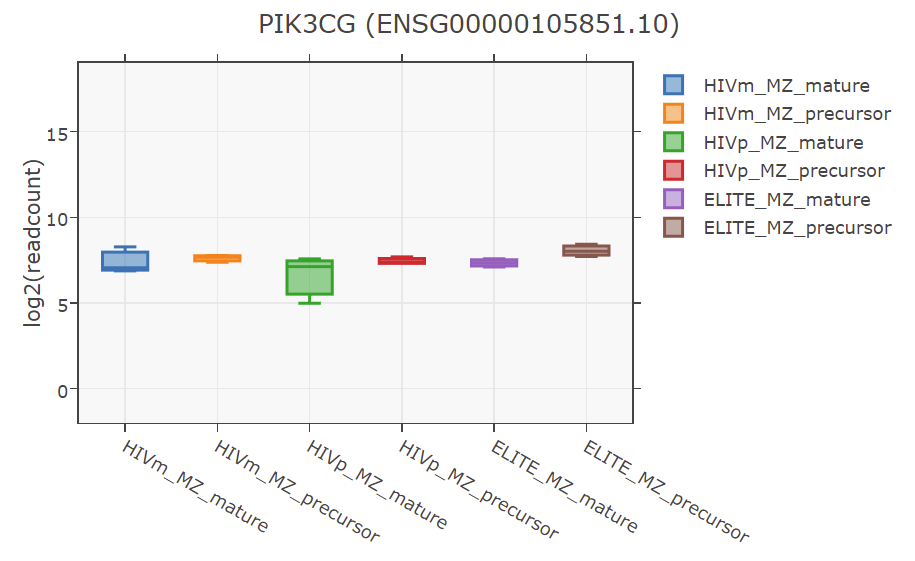

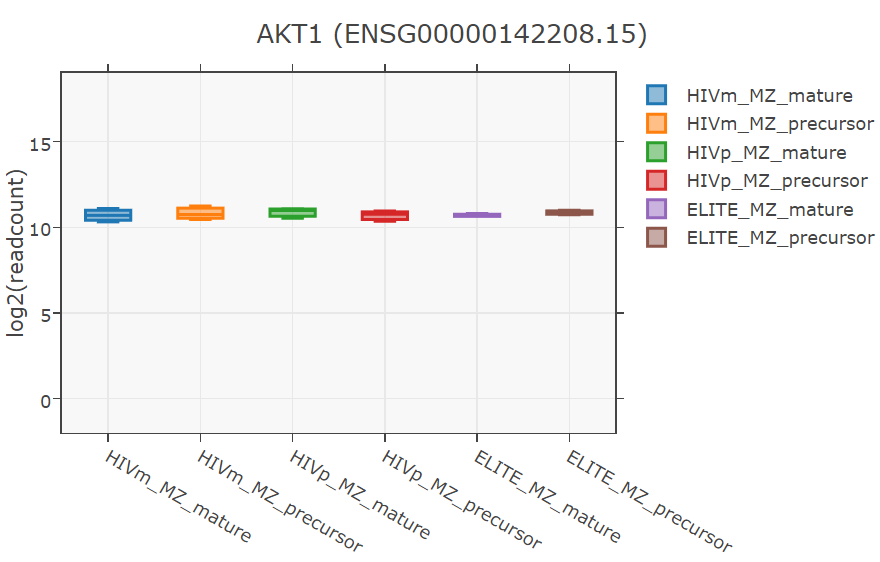

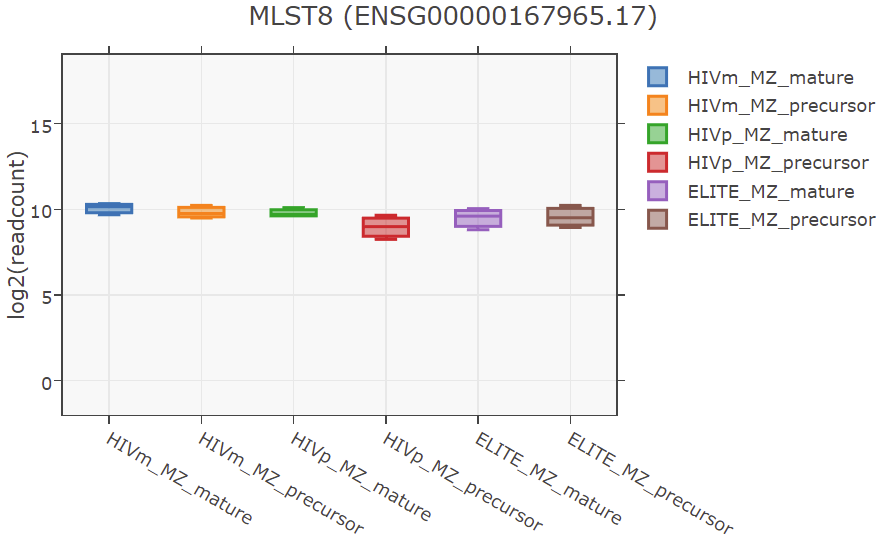

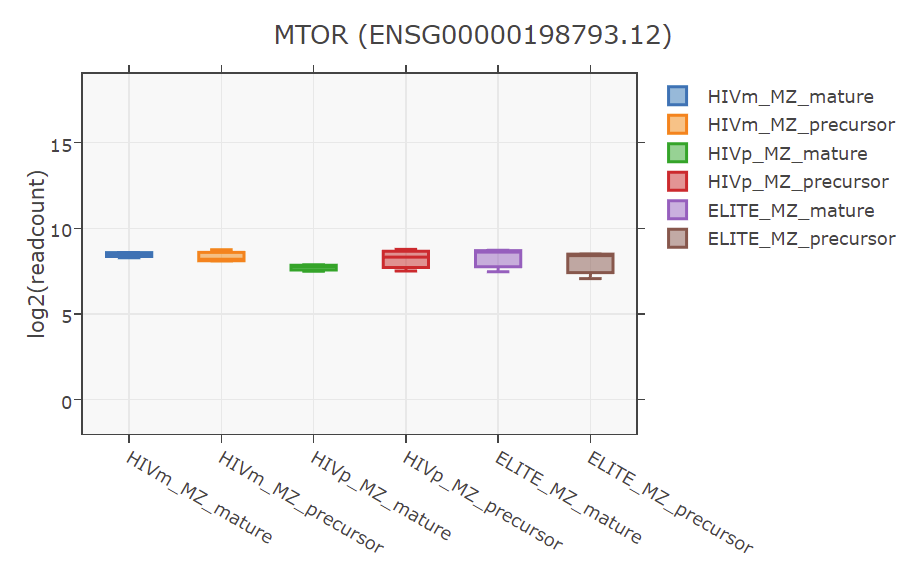

O

N

M

J

K

L

I

H

F

E

D

A

**mTOR**

**MLST8**

B

C

**0,09**
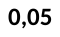

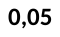

**S**

**NS**

**PIK3CG**

**AKT1S1**

**0,001**

**S**

**0,08**

**S**

**NS**

**S**

**0,02**

**S**

**0,003**

**S**

**NS**

**S**

**0,08**

**S**

**NS**

**S**

**NS**

**S**

**NS**

**S**

**NS**

**S**

**NS**

**S**

**0,05**

**S**

**PRKAR2A**

**PRKAR1A**

**PRKACA**

**Fos**

**Jun**

**AKT3**

**AKT2**

**AKT1**

**PIK3CD**

**RPTOR**

**DEPTOR**

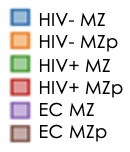
**Supplementary Figure 3. Blood MZp B-cells from HIV-infected progressors express lower levels of PD-L1 than uninfected controls.** Flow cytometry analyses of 1 uninfected control, 2 HIV-infected progressors and 3 ART-treated HIV-infected progressors. Shown are the relative frequencies of PD-L1 expressing total B-cells (**A**), MZ (**C**) and MZp (**E**) as well as the expression levels of PD-L1 by total B-cells (**B**), MZ (**D**) and MZp (**F**). Relative frequencies were assessed relatively to the percentage of total B-cells, MZ and MZp B-cells, respectively. Expression levels were assessed with Geometric Mean of Fluorescence Intensity (GeoMFI). Statistical differences between groups were assessed with the Kruskal-Wallis test with Dunn’s post-hoc test. Normality was assessed with the Shapiro-Wilk test.

G

A

B

C

D

E

F

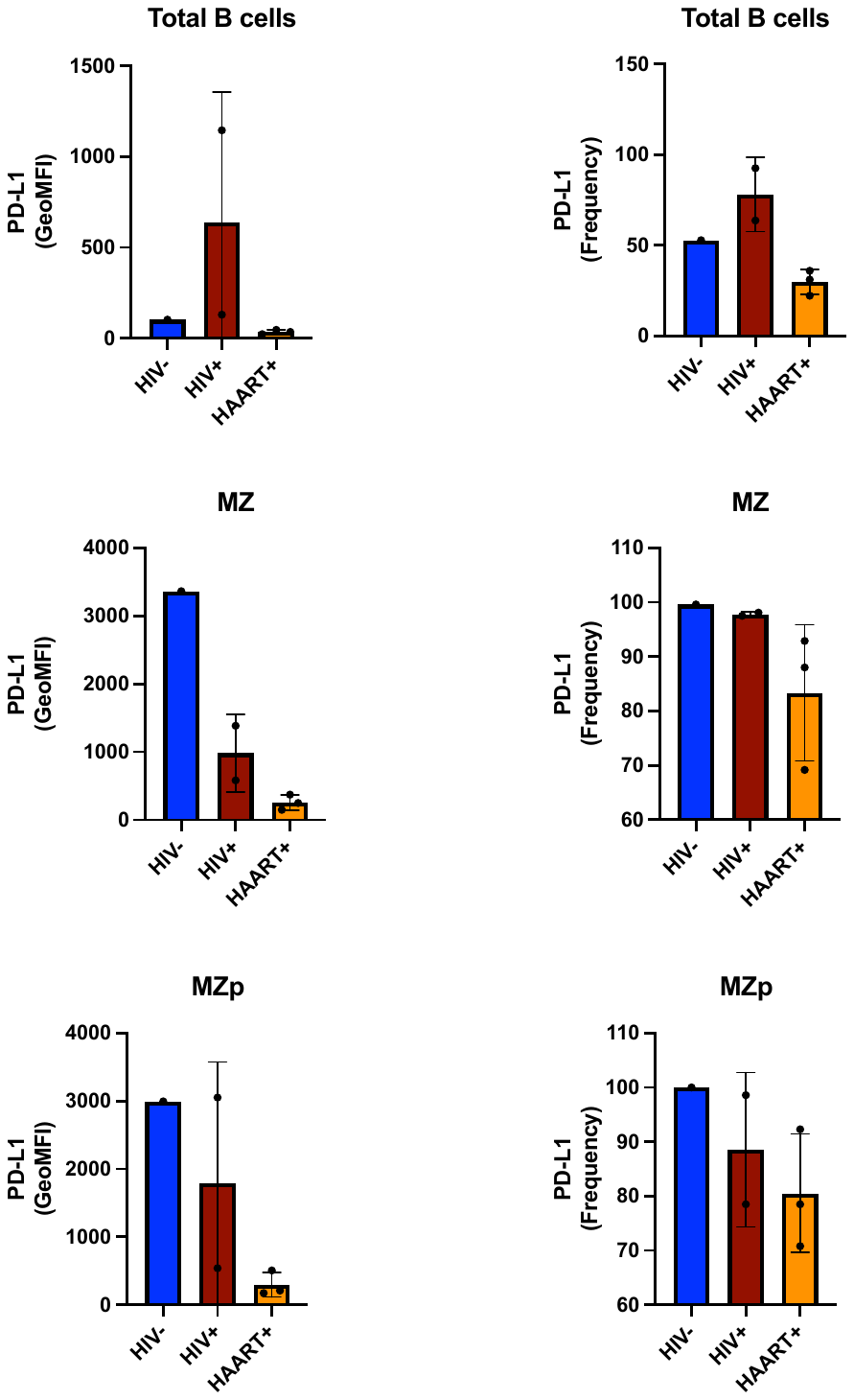

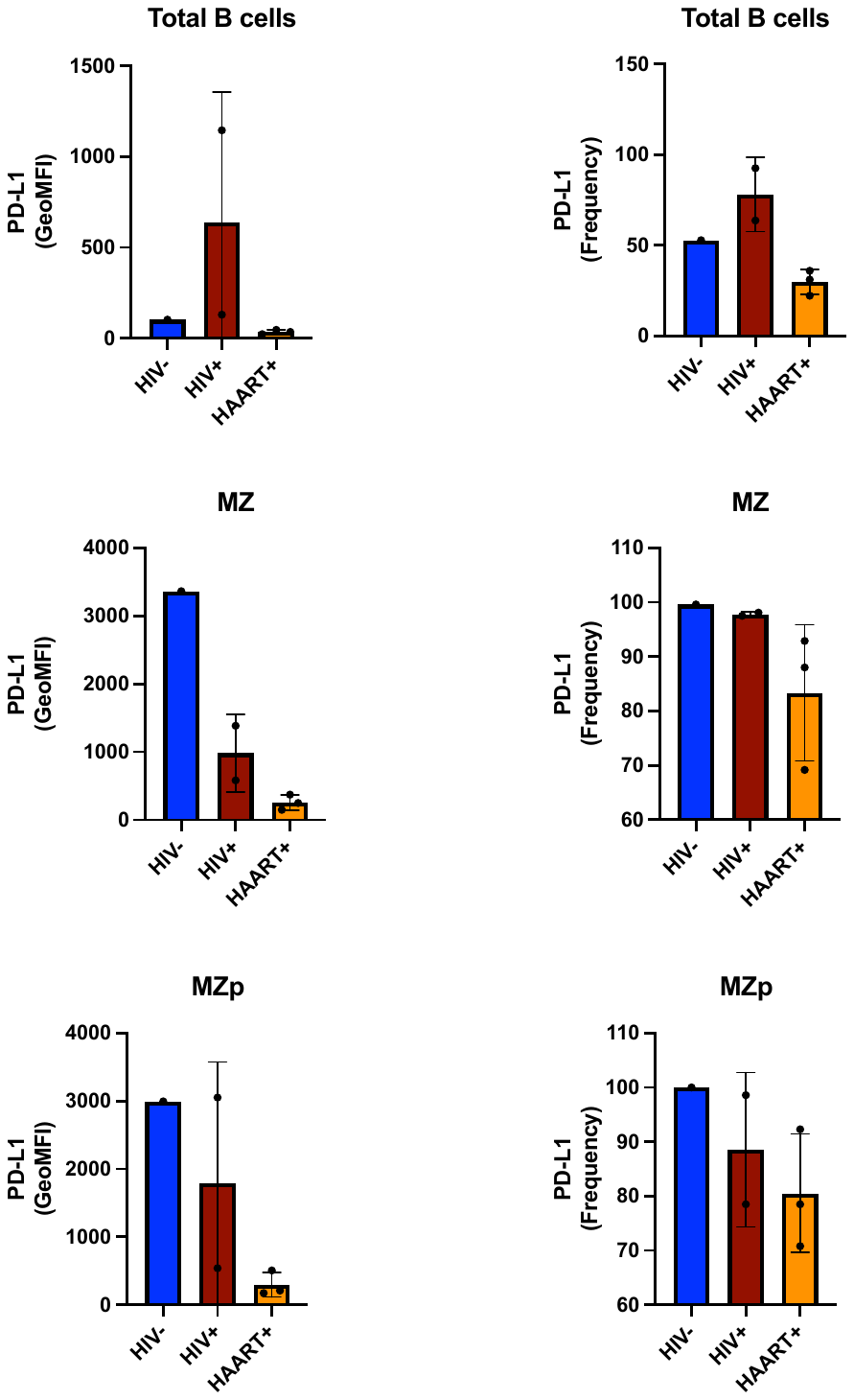

**Supplementary Figure 4. Data exploration of the transcriptomic analyses by RNAseq of** **blood MZp B-cells.** Data exploration of the transcriptomic analyses by RNAseq of sorted mature marginal zone (MZ) and MZ precursor (MZp) B-cells from the blood of 5-8 months HIV-infected progressors (HIV+), elite controllers (EC) and uninfected controls (HIV-) (n=3 for each group). Shown are gene expression levels of TLR7 (**A**), IL-21 (**B**), TLR10 (**C**), BAFFR (**D**), TACI (**E**), BCMA (**F**), TRAF3 (**G**), TRAF6 (**H**), IL-10 (**I**), AhR (**J**), BHLHE40 (**K**), CD38 (**L**). N = 3 for each group of participants. Statistical analyses were done between blood MZp B-cells from uninfected controls (HIV- MZp) and HIV-infected progressors (HIV+ MZp). The Wald Test with Benjamini-Hochberg correction was used for RNAseq analysis.

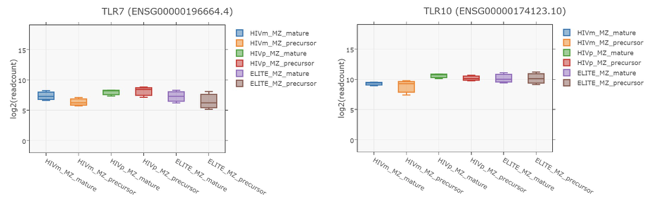

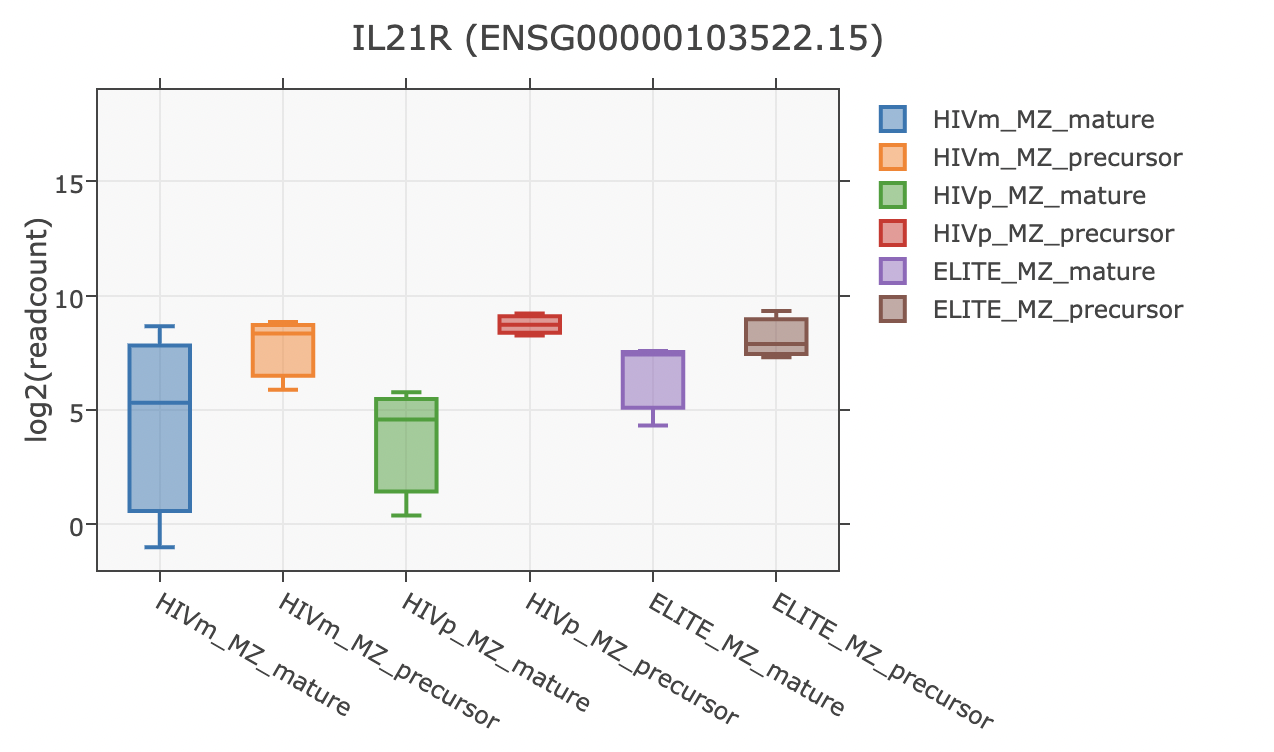

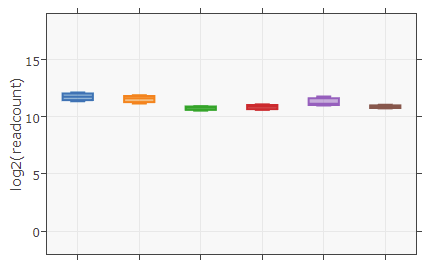

**TLR7**

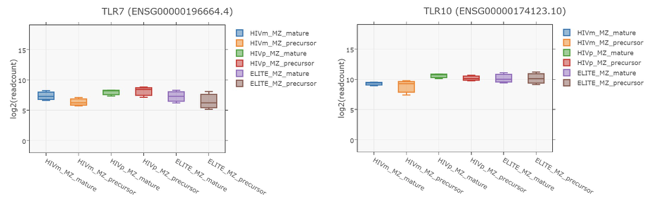

**IL-21**

**TLR10**

**BAFFR**

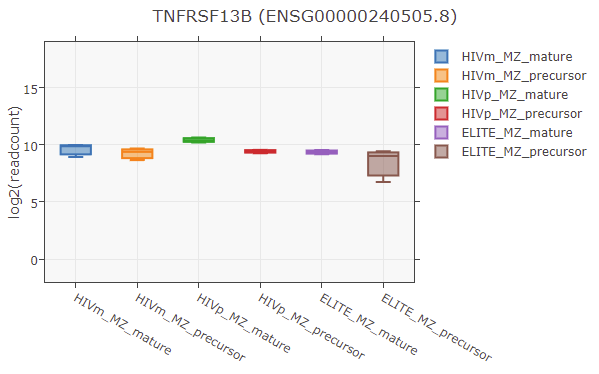

**TACI**

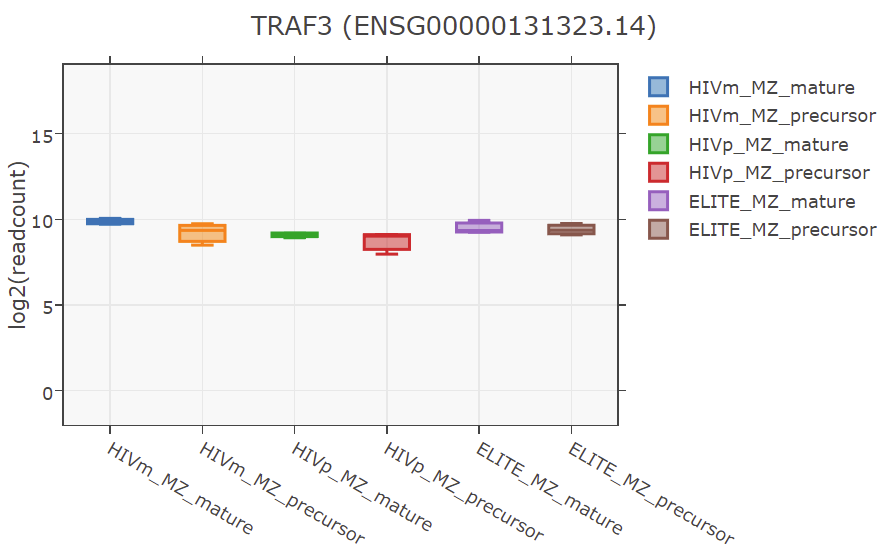

**TRAF3**

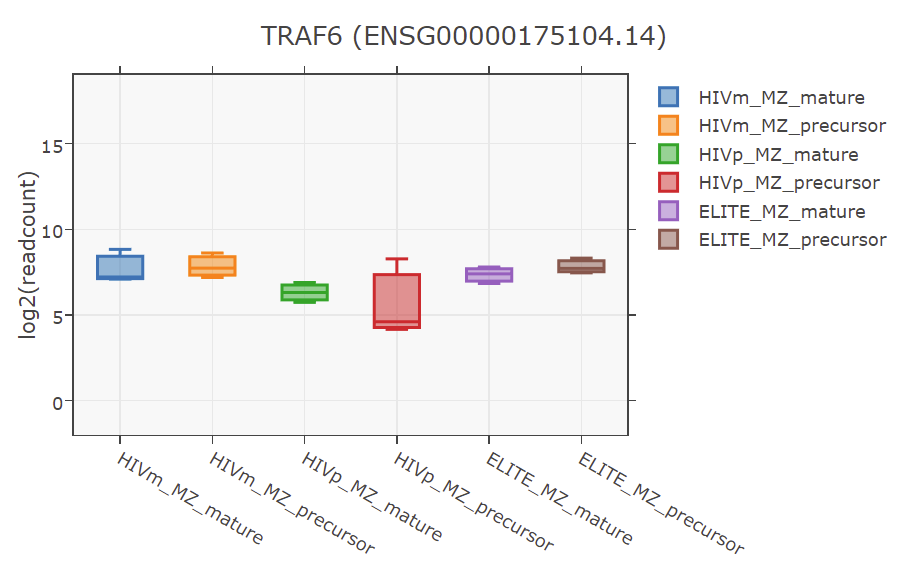

**TRAF6**

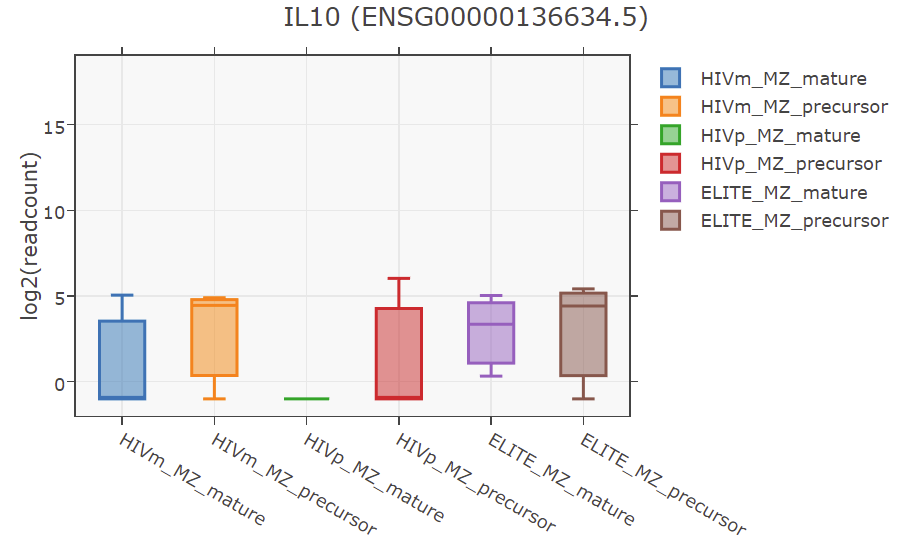

**IL-10**

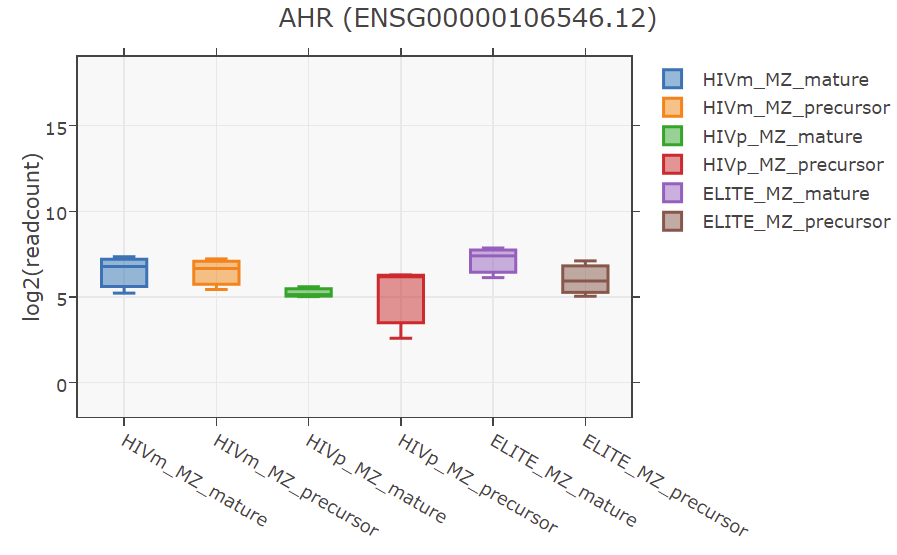

**AhR**

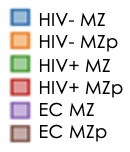

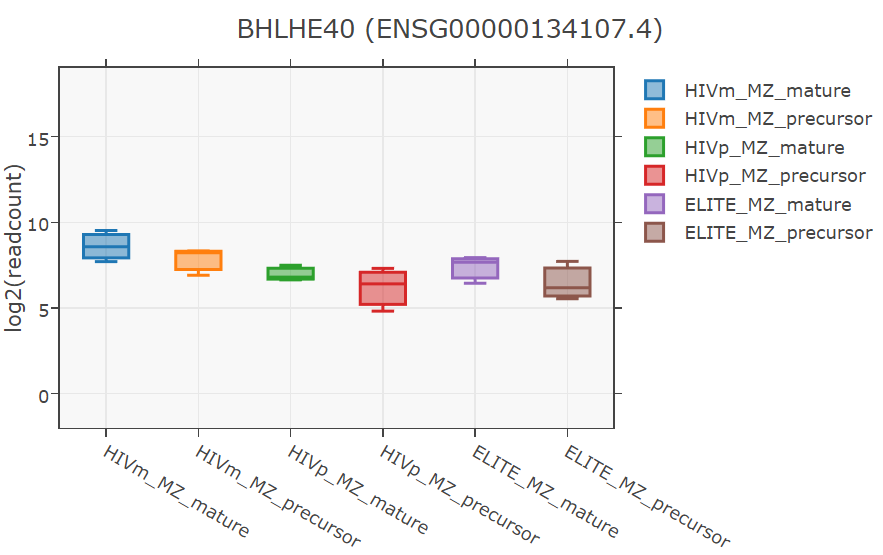

**BHLHE40**

A

B

C

D

E

F

G

H

I

J

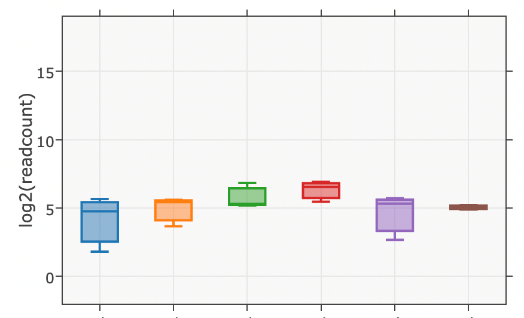

**BCMA**

**NS**

**0,05**

**0,01**

**NS**

**0,007**

**0,1**

**NS**

**NS**

**0,17**

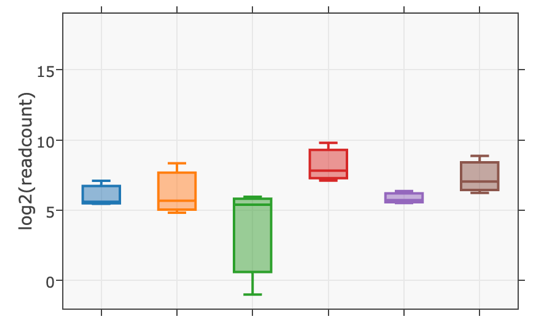

**CD38**

L

**0,08**

K

**0,03**

**NS**

**Supplementary Figure 5. Modulation of TACI and BAFF-R expression on B-cells in presence or absence of soluble BAFF.** (A) Dot plots show analysis of live B-cells for BAFF-R expression patterns, in medium alone (left upper panel) or in presence of BAFF (left lower panel). BAFF-R+ B-cells are further characterized upon their expression levels of CD27, CD21, CD10 and CD1c, in medium alone (upper panels) or in presence of BAFF (lower panels). (B) Dot plots show live B-cells analysis for TACI expression patterns, in medium alone (left upper panel) or in presence of BAFF (left lower panel). TACI+ IgM+ B-cells are further characterized upon their expression levels of CD19, CD21, CD27, CD10 and CD1c, in medium alone (upper panels) or in presence of BAFF (lower panels). Quadrants are set based on the expression values obtained with fluorescence minus one (FMO) and isotype controls.

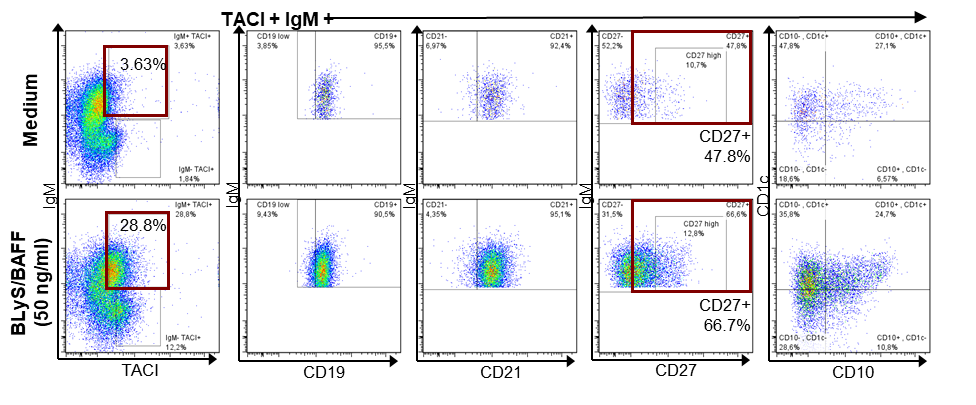

B

16,1%

6,82%

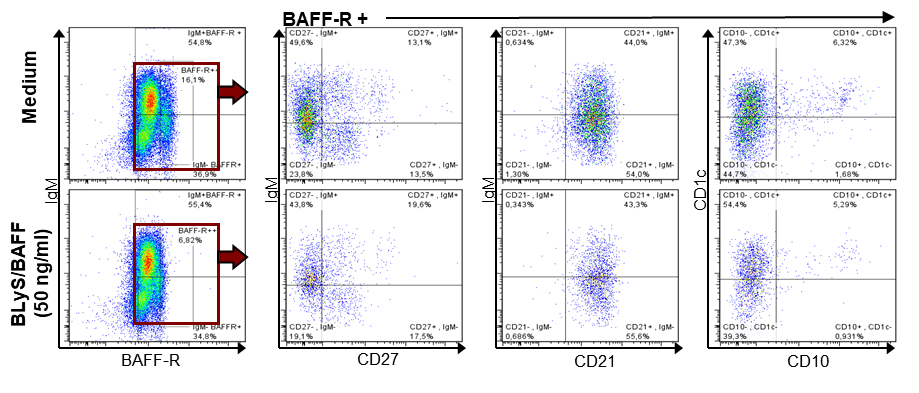

A

**Supplementary Figure 6. Tonsillar MZp B-cells express high levels of IL-10 without stimulation.** Flow cytometry analyses of intracellular IL-10 expression by MZp from one tonsillar donor (n=3) after 4 hours of Brefeldin A incubation. Shown are the relative frequencies of IL-10 expressing total B-cells, MZ and MZp (**A**), as well as the expression levels of IL-10 in total B-cells, MZ and MZp (**B**). Frequencies were assessed relatively to the percentage of total B-cells, MZ and MZp B-cells, respectively. Expression levels were assessed with Geometric Mean of Fluorescence Intensity (GeoMFI). *P < 0,05; ** P < 0,01; *** P < 0,001; **** P < 0,0001. Statistical differences between groups were assessed with the Kruskal-Wallis test with Dunn’s post-hoc test.

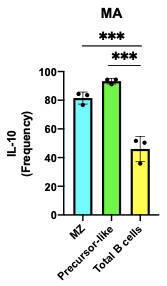

A

B

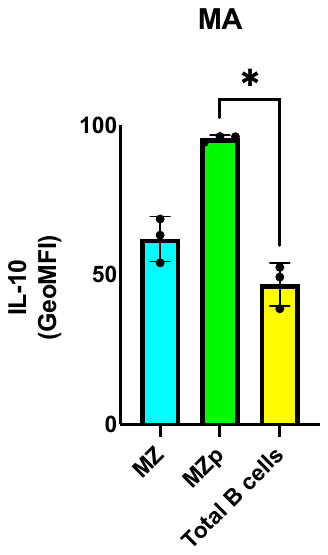

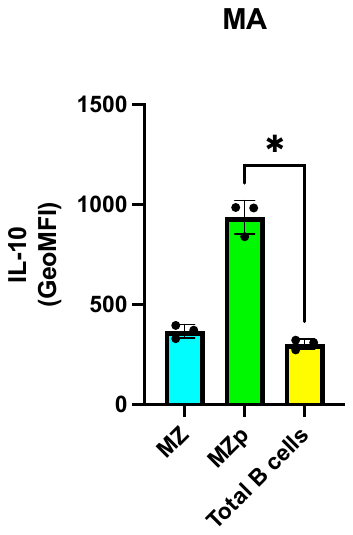

**Supplementary Figure 7. LPS stimulated MoDC downregulate NR4A1 and NR4A3 expression levels in autologous co-cultured blood MZp B-cells from healthy individuals.** Flow cytometry analyses of MZp B-cells cultured in medium alone (MA) or with autologous MoDC, which had been stimulated with LPS and fixed prior to co-culture (MoDC LPS). Shown are the relative frequencies of NR4A1 (**A**) and NR4A3 (**C**) expressing MZp B-cells as well as expression levels of NR4A1 (**B**), NR4A3 (**D**), and membrane BAFF expression levels by MoDC and MoDC LPS (**E**). Frequencies were assessed relatively to the percentage of total MZp B-cells. Expression levels were assessed with Geometric Mean of Fluorescence Intensity (GeoMFI). (n=2). MA - Medium Alone, MoDC - Monocyte Derived Dendritic Cells; LPS - Lipopolysaccharide. Statistical differences between groups were assessed with the Kruskal-Wallis test with Dunn’s post-hoc test (Supp. Fig. 6 A-D).

A

B

C

D

E

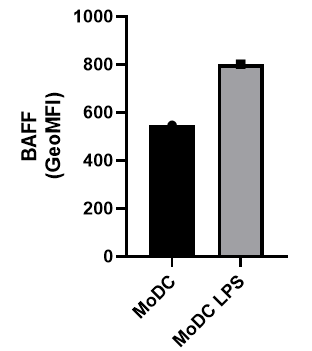

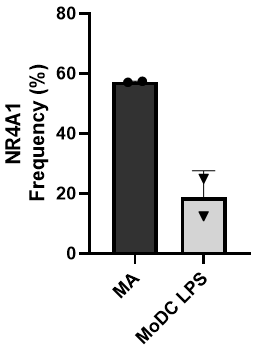

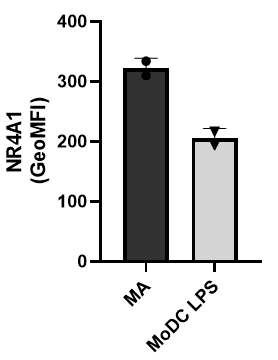

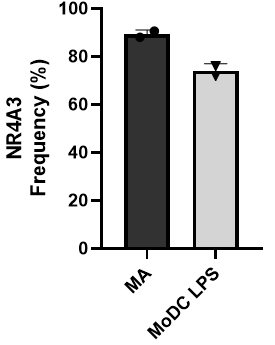

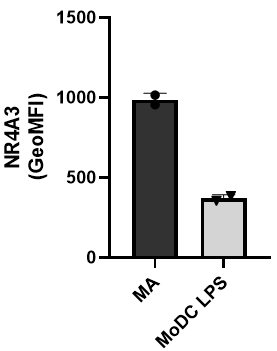
